# Jag1 modulates an oscillatory Dll1-Notch-Hes1 signaling module to coordinate growth and fate of pancreatic progenitors

**DOI:** 10.1101/336529

**Authors:** Philip A. Seymour, Caitlin A. Collin, Anuska l. R. Egeskov-Madsen, Mette C. Jørgensen, Hiromi Shimojo, Itaru Imayoshi, Kristian H. de Lichtenberg, Raphael Kopan, Ryoichiro Kageyama, Palle Serup

**Affiliations:** Novo Nordisk Foundation Center for Stem Cell Biology (DanStem), University of Copenhagen, Blegdamsvej 3B, DK-2200, Copenhagen N, Denmark; Institute for Frontier Life and Medical Sciences, Kyoto University, Kyoto, 606-8507, Japan; Division of Developmental Biology, Cincinnati Children’s Hospital Medical Center, Cincinnati, OH 45229, USA

**Keywords:** Notch, oscillations, cis-inhibition, pancreas, development, fate, Dll1, Jag1, Hes1

## Abstract

Notch signaling controls proliferation of multipotent pancreatic progenitor cells (MPCs) and their segregation into <u>b</u>ipotent <u>p</u>rogenitors (BPs) and unipotent <u>p</u>ro-<u>ac</u>inar cells (PACs). Here we uncover fast ultradian oscillations in the ligand Dll1, and the transcriptional effector Hes1, which proved crucial for MPC expansion. Conversely Jag1, a uniformly expressed ligand, curbed MPC growth, but as expression later segregated to PACs it proved critical for specifying all but the most proximal 5% of BPs, while BPs were entirely lost in *Jag1*, *Dll1* double mutants. Moreover, experimentally induced changes in Hes1 oscillation parameters was associated with selective adoption of BP or PAC fates. Anatomically, ductal morphogenesis and organ architecture is minimally perturbed in *Jag1* mutants until later stages, when ductal remodeling fails and signs of acinar-to-ductal metaplasia appear. Our study uncovers oscillating Notch activity in the developing pancreas, which along with modulation by Jag1 is required to coordinate MPC growth and fate.

## Introduction

Deciphering the mechanisms that control the differentiation of progenitor cells into various mature cell types is crucial for understanding disease etiology and for using pluripotent stem cells in cell replacement therapy applications. The principal cell lineages of the mammalian pancreas, acinar, duct and endocrine, arise from multipotent pancreatic progenitor cells (MPCs) through a series of binary cell fate choices (Shih et al., 2013). MPCs are specified from the posterior foregut endoderm as dorsal and ventral anlagen around embryonic day (E)8.5 (Wessells and Cohen, 1967), and are distinguished by their expression of several transcription factors including Pdx1, Ptf1a, Hnf1β, Sox9 and Nkx6-1. The multipotentiality of these cells has been demonstrated at the population level by many lineage-tracing studies (Gu et al., 2002; Kopp et al., 2011; Pan et al., 2013; Solar et al., 2009; Zhou et al., 2007) and, more recently, at the clonal level (Larsen et al., 2017).

Dorsal MPCs give rise to an early wave of endocrine cells from ~E9.5 to ~E11 while endocrinogenesis is delayed ~36 hours in the ventral anlage (Ahlgren et al., 1997; Pictet et al., 1972; Spooner et al., 1970). The appearance of hormone-producing cells is preceded by, and dependent on, expression of *Neurog3* (encoding Ngn3) in endocrine precursors, which initiates at ~E8.5 in the dorsal bud and ~E10.0 in the ventral bud (Gradwohl et al., 2000; Gu et al., 2002; Villasenor et al., 2008). A transient decline in *Neurog3* expression from E11 to E12 (Villasenor et al., 2008) coincide with peak segregation of Ptf1a^+^Nkx6-1^+^ MPCs into proximal, Ptf1a^−^Nkx6-1^+^ bipotent progenitors (BPs) and distal, Ptf1a^+^Nkx6-1^−^ pro-acinar cells (PACs) (Hald et al., 2008; Schaffer et al., 2010). This proximodistal (PD) patterning is regulated by Notch signaling as forced expression of a Notch intracellular domain (NICD) prevents MPCs from adopting a PAC fate (Hald et al., 2003; Murtaugh et al., 2003; Schaffer et al., 2010) while dominant-negative Maml1 prevents a BP fate (Afelik et al., 2012; Horn et al., 2012). Total inactivation of Notch signaling in the endoderm, as seen in *Foxa2*^T2AiCre/+^; *Mib1*^f/f^ (*Mib1*^ΔFoxa2^) embryos, results in a complete shift from BP- to PAC fate (Horn et al., 2012). Downstream of Notch, PD patterning depends on mutually antagonistic interactions between Ptf1a and Nkx6-1/Nkx6-2(Schaffer et al., 2010). Notch, and its downstream target gene *Hes1*, also prevent precocious and excessive endocrine differentiation (Apelqvist et al., 1999; Jensen et al., 2000b) and stimulate MPC proliferation (Ahnfelt-Rønne et al., 2012). After PD patterning is complete around E14, Notch signaling and Hes1 expression persist in BPs, maintaining Sox9 expression and ductal fate, and inhibiting endocrine differentiation by repressing *Neurog3* (Bankaitis et al., 2015; Klinck et al., 2011; Kopinke et al., 2011; Magenheim et al., 2011; Shih et al., 2012). Less is known about which ligands regulate the adoption of distinct cell fates. Dll1 regulates early endocrine differentiation (Ahnfelt-Rønne et al., 2012; Apelqvist et al., 1999), yet PD patterning appears unaffected in *Dll1*^ΔFoxa2^ embryos and the endocrinogenic phenotype of E10.5 *Dll1*^ΔFoxa2^ embryos is much weaker than that of E10.5 *Mib1*^ΔFoxa2^ embryos (Horn et al., 2012; Jorgensen et al., 2018). These findings show that additional Mib1 substrates, most likely other Notch ligands, are involved in these cell fate decisions. In zebrafish, intrapancreatic duct cells are depleted in *jag1b*/*jag2b* morphants (Lorent et al., 2004; Yee et al., 2005) and completely absent from *jag1b*^−/−^; *jag2b*^−/−^ embryos (Zhang et al., 2017) with the latter also showing a reduction in endocrine cells while acinar cells were unaffected. However, in mice Jag1 is the only Jagged-type ligand expressed in the pancreatic endoderm (Lammert et al., 2000), and *Jag1*^ΔPdx1^ animals displayed ductal malformation and paucity postnatally, due to reduced duct cell proliferation (Golson et al., 2009b). Similarly, *Dll1*; *Jag1*^ΔPtf1a^ embryos showed a modest phenotype with loss of only the terminal duct or centroacinar cells (CACs) (Nakano et al., 2015). Thus, which ligands that regulate the various cell fate decisions remains unclear as do dynamics and timing of Notch signaling. In this study, we uncover a complex interaction between Dll1 and Jag1 that regulates growth and differentiation of pancreatic progenitors. We demonstrate that Dll1 and Hes1 expression oscillates and that oscillations are important for progenitor expansion and fate choice. Conversely, we find uniformly expressed Jag1 to attenuate Notch activity cell-autonomously. Yet, Jag1 can activate Notch in Jag1^−^ neighbors, in partial redundancy with Dll1, and together the two ligands specify the entire BP lineage. Anatomically, epithelial plexus formation and the gross architecture of the organ are remarkably unperturbed in spite of the profound changes in cell fate. However, at late stages larger ducts are missing or malformed, terminal ducts are morphologically abnormal and signs of acinar-to-ductal metaplasia (ADM) appear in *Jag1*^ΔFoxa2^ embryos.

## Results

### Initially uniform Jag1 expression segregates to nascent PACs

Since BPs are converted to PACs in *Mib1*^ΔFoxa2^ embryos, but not in *Dll1*^ΔFoxa2^ embryos (Horn et al., 2012), we reasoned that an additional Mib1 substrate, most likely Jag1 (Ahnfelt-Rønne et al., 2012; Apelqvist et al., 1999; Golson et al., 2009a; Jensen et al., 2000a; Lammert et al., 2000; Nakano et al., 2015), must act during PD patterning. Because the reported expression patterns are not entirely consistent, we decided to re-analyze the expression of these two ligands using fluorescent protein reporters targeted to the *Dll1* and *Jag1* loci. The *Jag1*^J1VmC^ and *Dll1*^D1VmC^ alleles comprise a Venus-T2A-mCherry cassette inserted in frame with the coding region of the last exons to generate Dll1- and Jag1-Venus fusion proteins that serve as dynamic reporters of ligand protein expression while mCherry acts as a more sensitive reporter owing to its longer half-life after being cleaved from the fusion protein. Co-immunofluorescence (IF) analysis with anti-GFP antibodies revealed uniform Jag1-Venus fusion protein expression in E10.5 *Jag1*^J1VmC/+^ Pdx1^+^Sox9^+^ MPCs in the dorsal pancreas epithelium and weaker expression in the surrounding mesenchyme and a restriction to the distal epithelium at E12.5, with most of the centrally located, emerging Sox9^+^ BPs being negative for Jag1-Venus. At E15.5, Jag1-Venus expression was detected apically in acinar cells and in non-parenchymal cells, including the vasculature, but expression was excluded from Sox9^+^ BPs and Pdx1^Hi^ β-cells (Figure 1A). In *Dll1*^D1VmC^ embryos we detected Dll1-Venus fusion protein in scattered cells throughout the E10.5 and E12.5 dorsal pancreatic epithelium, some of which were Sox9^Lo/−^, and by E15.5, Dll1-Venus was found intracellularly in the apical pole of acinar cells, and in dispersed Sox9^Lo/−^ cells in the proximal epithelium that likely represent endocrine precursors (Figure 1A). Anti-RFP antibodies confirmed these expression patterns and further revealed onset of Jag1 expression in MPCs between E9.5 and E10.5 (Figure S1A). Similarly, validated anti-Jag1 and anti-Dll1 antibodies (see Methods, Figure S7A-7L) reproduced the expression patterns of the Jag1- and Dll1-reporters in wild type pancreas, and IF analysis of lineage markers confirmed Jag1 expression in vascular cells and acinar cells at E15.5, and its absence from E15.5 Sox9^+^ BPs and from the endocrine lineage at all stages (Figure S1B). Equally, Dll1^+^ cells could be divided into subpopulations comprising Dll1^+^Ptf1a^+^ MPCs or PACs present in E10.5, E12.5, and E15.5 pancreas epithelium as well as Dll1^+^Ngn3^+^ endocrine precursors, evident at all stages, and Dll1^+^Ngn3^−^ cells in E12.5 proximal epithelium (Figure S1C). Lastly, co-expression of Dll1 and Jag1 was evident in some E10.5 MPCs and in a subset of distal cells at E12.5 (Figures 1B and 1C). Co-IF for Jag1, Dll1, Ptf1a and Nkx6-1 revealed that these were Ptf1a^+^Nkx6-1^Lo/−^ nascent PACs and MPCs and in the proximal region Dll1^+^ cells were Ptf1a^−^Jag1^−^ and either Nkx6-1^+^ or Nkx6-1^−^ (Figure 1C).

**Figure 1.**
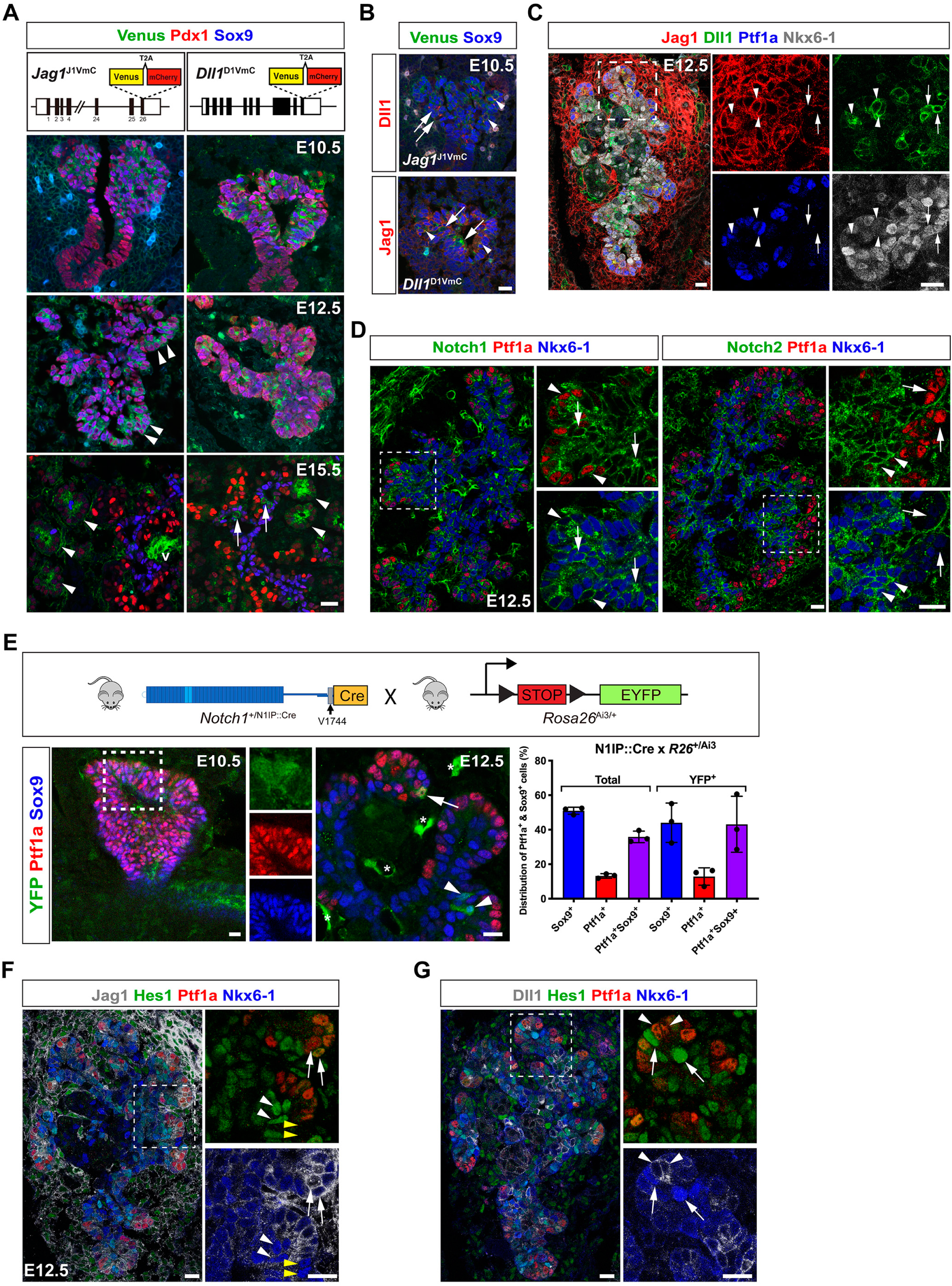
Differential expression of Notch components in MPCs and their progeny. (A) Sections of E10.5, E12.5 and E15.5 *Jag1*^J1VmC^ and *Dll1*^D1VmC^ dorsal pancreata stained for Venus, Pdx1 and Sox9 as indicated. Arrowheads in E12.5 panel indicate emerging PACs. Arrows in E15.5 panels indicate endocrine precursors and arrowheads indicate forming acini. v: vessel. Lower panels show schematic representations of the *Jag1*^J1VmC^ and *Dll1*^D1VmC^ fusion protein reporters. (B) Sections of E10.5 *Jag1*^J1VmC^ and *Dll1*^D1VmC^ dorsal buds stained for Venus, Sox9 and Dll1 or Jag1 as indicated. Arrows indicate Dll1^+^Jag1^−^ cells and arrowheads indicate Sox9^+^ MPCs co-expressing Jag1 and Dll1. (C) Section of E12.5 dorsal pancreas stained for Ptf1a, Nkx6-1, Dll1 and Jag1 as indicated. Arrows indicate Nkx6-1^+^ BPs expressing Dll1 but not Jag1 and arrowheads indicate Ptf1a^+^ PACs co-expressing Jag1 and Dll1. (D) Serial sections of E12.5 dorsal pancreas stained for Ptf1a, Nkx6-1, and Notch1 or Notch2 as indicated. Arrows and arrowheads in Notch1 panels indicate Notch1^+^Nkx6-1^+^ and Notch1^+^Ptf1a^+^ cells, respectively. Arrows and arrowheads in Notch2 panels indicate Notch2^−^Ptf1a^+^ and Notch2^+^Nkx6-1^+^ cells, respectively. (E) Upper panel shows a schematic of the lineage-tracing strategy using N1IP::Cre to label cells with a history of Notch1 activation. Lower panels show sections of E10.5 and E12.5 dorsal pancreas from an N1IP::Cre; Rosa26LSL-Ai3 embryo stained for YFP, Ptf1a and Sox9 as indicated. Insets show individual channels for clarity. Arrow: YFP-labeled, Ptf1a^+^ PAC. Arrowheads: YFP-labeled, Sox9^+^ BPs. Asterisks: YFP-labeled endothelial cells. The bar graph shows quantification of YFP-labeled versus unlabeled epithelial cells expressing Sox9, Ptf1a, or both. Mean ± S.D., N = 3 embryos. (F) Section of E12.5 dorsal pancreas stained for Ptf1a, Nkx6-1, Hes1 and Jag1 as indicated. Arrows: Ptf1a^+^Jag1^+^Nkx6-1^Lo/−^ cells. Arrowheads: Nkx6-1^+^ cells; either Hes1^+^ (white arrowheads) or Hes1^−^ (yellow arrowheads). (G) Section of E12.5 dorsal pancreas stained for Ptf1a, Nkx6-1, Hes1 and Dll1 as indicated. Arrows: Nkx6-1^Hi^Hes1^Hi^ cells in distal epithelium adjacent to Ptf1a^+^Dll1^Hi^ cells (arrowheads). Scale bars are 20 μm in all panels. See also Figures S1 and S7.

### Notch activity is suppressed in nascent PACs

We next examined the distribution of Notch1 and Notch2 in emerging BPs and PACs of the E12.5 pancreatic epithelium. While Notch1 was expressed in PACs and BPs, Notch2 was specifically enriched in BPs (Figure 1D), as previously noted in E15.5 pancreas (Shih et al., 2012). To identify cells in which Notch1 had been activated before and during PD patterning we first analyzed embryonic pancreata from the Notch1 activity-trap mouse line N1IP::Cre^HI^; *Rosa26*^LSL-Ai3^ in which EYFP permanently labels the progeny of cells experiencing Notch1 activation (Liu et al., 2015). We detected YFP labeling in a few E10.5 Ptf1a^+^Sox9^+^ MPCs and at E12.5 we found YFP expression in Ptf1a^+^ nascent PACs, Sox9^+^ nascent BPs and in Ptf1a^+^Sox9^+^ MPCs (Figure 1E). Quantification revealed that YFP^+^ cell distribution reflected no bias towards either nascent BPs or PACs at E12.5, consistent with the cells experiencing Notch1 activation in MPCs, from which both lineages derive, prior to the linage segregation occurring by E12.5. We next analyzed Hes1 expression as an acute readout of general, pan-Notch activation in relation to ligand protein expression in E12.5 MPCs and emerging Nkx6-1^+^ BPs and Ptf1a^+^ PACs. Most Hes1^Hi^ cells were Nkx6-1^Hi^Ptf1a^Lo/−^Jag1^Lo/−^ BPs and most Hes1^Lo/−^ cells were Nkx6-1^Lo/−^Ptf1a^Hi^Jag1^Hi^ PACs except for a few Hes1^Lo/−^ cells that were Nkx6-1^Hi^Ptf1a^Hi^Jag1^Hi^ (Figure 1F). In relation to Dll1, we found that Hes1^Hi^ cells typically were Nkx6-1^Hi^Ptf1a^Lo/−^Dll1^Lo/−^ BPs and, when located in the periphery, often located next to Hes1^Lo^Nkx6-1^Lo/−^Ptf1a^Hi^Dll1^Hi^ cells (Figure 1G). Analyses of double reporter mice harboring either *Jag1*^J1VmC^ or *Dll1*^D1VmC^ lines combined with a *Hes1-*EGFP reporter (Klinck et al., 2011) for EGFP and mCherry expression confirmed the progressive confinement of Hes1 to BPs and Jag1 to PACs as well as the heterogenous Dll1 expression at multiple stages, albeit the long half-life of EGFP and mCherry likely overestimated the number of co-positive cells (Figure S1D). A schematic representation of ligand and receptor expression patterns in shown in Figure S1E. Overall, we observe an inverse correlation between Dll1 and Hes1 expression and the data suggest that BPs (Jag1^−^Dll1^+/−^Notch1^+^Notch2^+^) have receptors in stoichiometric surplus, favoring signal reception, while ligands are in surplus in PACs (Jag1^+^Dll1^+/−^Notch1^+^Notch2^−^), favoring signal sending. Together, the data show that Notch receptor activation becomes suppressed in emerging Jag1^+^Dll1^+/−^ PACs, and support the notion that these, together with Ngn3^+^Dll1^Hi^ endocrine precursors in the central trunk epithelium, are activating Notch receptors in nascent, Jag1^Lo/−^Dll1^Lo/−^ BPs.

### Hes1 and Dll11 proteins display ultradian oscillations

An inverse correlation between heterogenous Dll1 and Hes1 expression is also observed in neural progenitors, where it reflects oscillating protein levels (Imayoshi et al., 2013; Shimojo et al., 2016; Shimojo et al., 2008), but it is unknown whether Notch components oscillate in the fetal pancreas. To address this question, we examined the dynamics of Hes1 and Dll1 protein expression in the developing pancreas using BAC-transgenic Luc2-Hes1 fusion protein reporter mice (Imayoshi et al., 2013) and *Dll1*-Fluc knock-in mice (Shimojo et al., 2016). E10.5 and E12.5 dorsal pancreata were explanted and cultured with luciferin, and bioluminescence images were examined. Time-lapse imaging analysis showed that Hes1 protein expression changed dynamically throughout the explant and neighboring cells were frequently observed to oscillate in anti-phase (Figure 2A, Movie S1-S3). Temporal analysis revealed well-defined oscillations in Hes1 protein levels with an average period of ~90 min that remained constant throughout a 6-day culture period (Figure 2A). Equally, time-lapse imaging analysis showed that Dll1 protein expression also changed dynamically throughout the explant (Figure 2B, Movie S4). Temporal analysis revealed that Dll1 protein levels also oscillated with an average period of ~90 min (Figure 2B). As both *Hes1* and *Dll1* are direct Hes1 target genes in pancreatic progenitors (de Lichtenberg et al., 2018a; de Lichtenberg et al., 2018b), these data indicate that Hes1 and Dll1 participate in the same oscillating gene regulatory network in the early pancreas.

**Figure 2.**
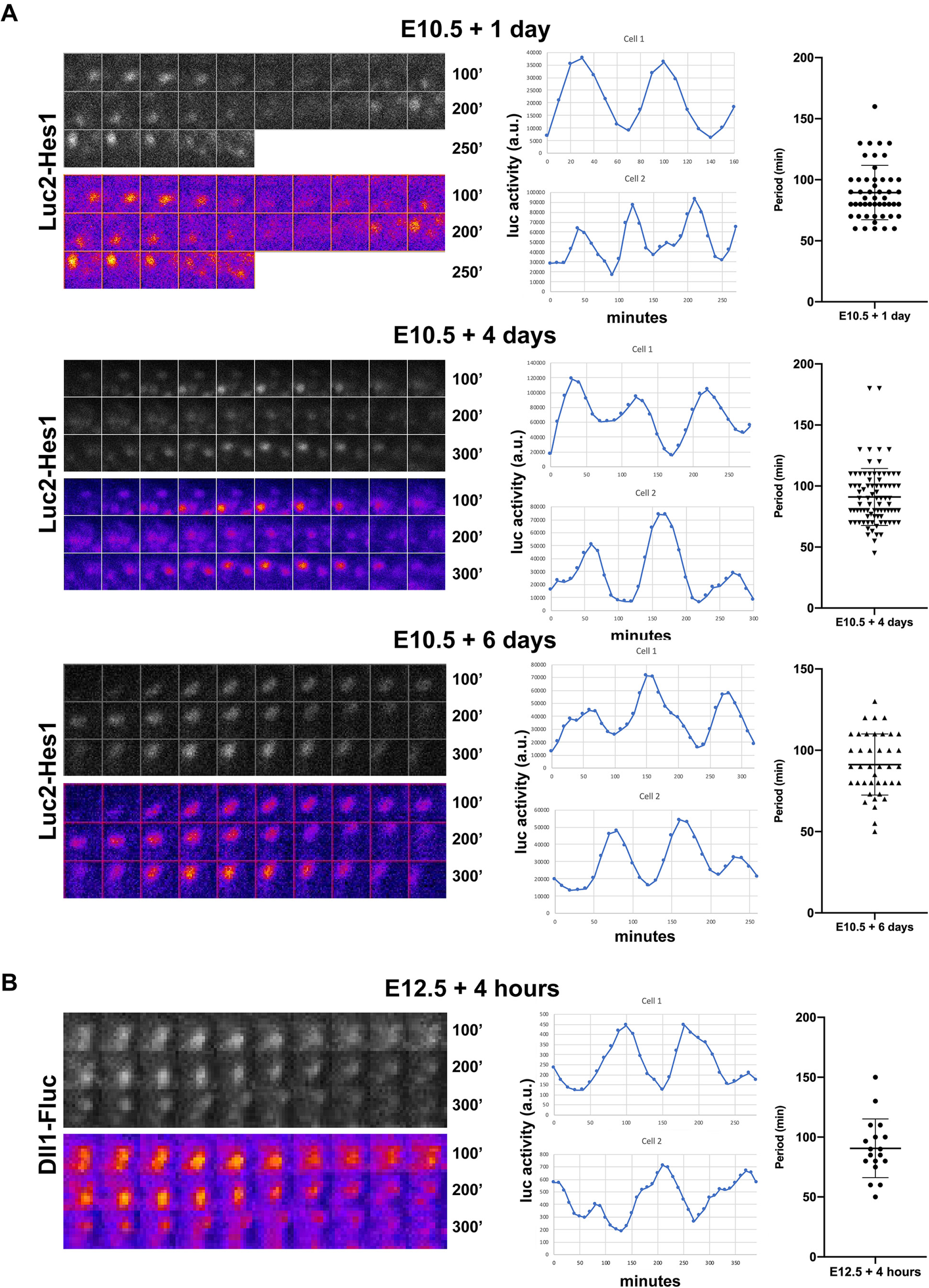
Hes1 and Dll1 protein levels display ultradian oscillations. (A) Bioluminescence images and quantification of Luc2-Hes1 reporter expression in E10.5 pancreatic explants cultured for 1, 4 or 6 days prior to time-lapse imaging. (B) Bioluminescence images and quantification of Dll1-Fluc reporter expression in E12.5 pancreatic explants cultured for 4 hours prior to time-lapse imaging. (A, B) Grayscale and false color montages are shown for the same cell and each frame is 10 minutes (10’). Representative tracks from two separate cells are shown for each condition and the scatter plots shows the distribution and average of the oscillation periods (Mean ± S.D., N > 50 cells for each Hes1-Luc2 time point and N = 18 for Dll1-Fluc).

### Dll1 oscillations augment and Jag1 attenuates MPC growth

We have previously shown that global mutations in either Hes1 or Dll1 reduces MPC proliferation and the size of the pancreas anlage (Ahnfelt-Rønne et al., 2012). Given the different spatiotemporal expression patterns of Dll1 and Jag1 in MPCs we first asked whether Dll1 oscillations were important for Dll1 stimulated MPC expansion. To address this question, we examined pancreatic bud size by whole-mount IF (WM-IF) in two different *Dll1* mutant mouse lines (termed *Dll1 type1* and *Dll1 type2* mutants) that display steady, intermediate level Dll1 expression and dampened Hes1 oscillations (Shimojo et al., 2016) and compared to littermate wild-type controls. We found that average dorsal pancreatic bud size was reduced by ~45% and ~25% in *Dll1*^type1/type1^ and *Dll1*^type2/type2^ mutants, respectively, while ventral buds were reduced by ~55% and ~65%, respectively (Figure 3A). The magnitude of this change is comparable to a ~65% reduction in dorsal and ventral bud size observed in E10.5 *Dll1*^ΔFoxa2^ embryos (Figure 3A), and in *Dll1*^−/−^ mutants (Ahnfelt-Rønne et al., 2012). These results suggest that Dll1 oscillations are important for MPC expansion by ensuring adequate Notch activation, perhaps by allowing MPCs to alternate between sending and receiving states multiple times within a single cell cycle.

**Figure 3.**
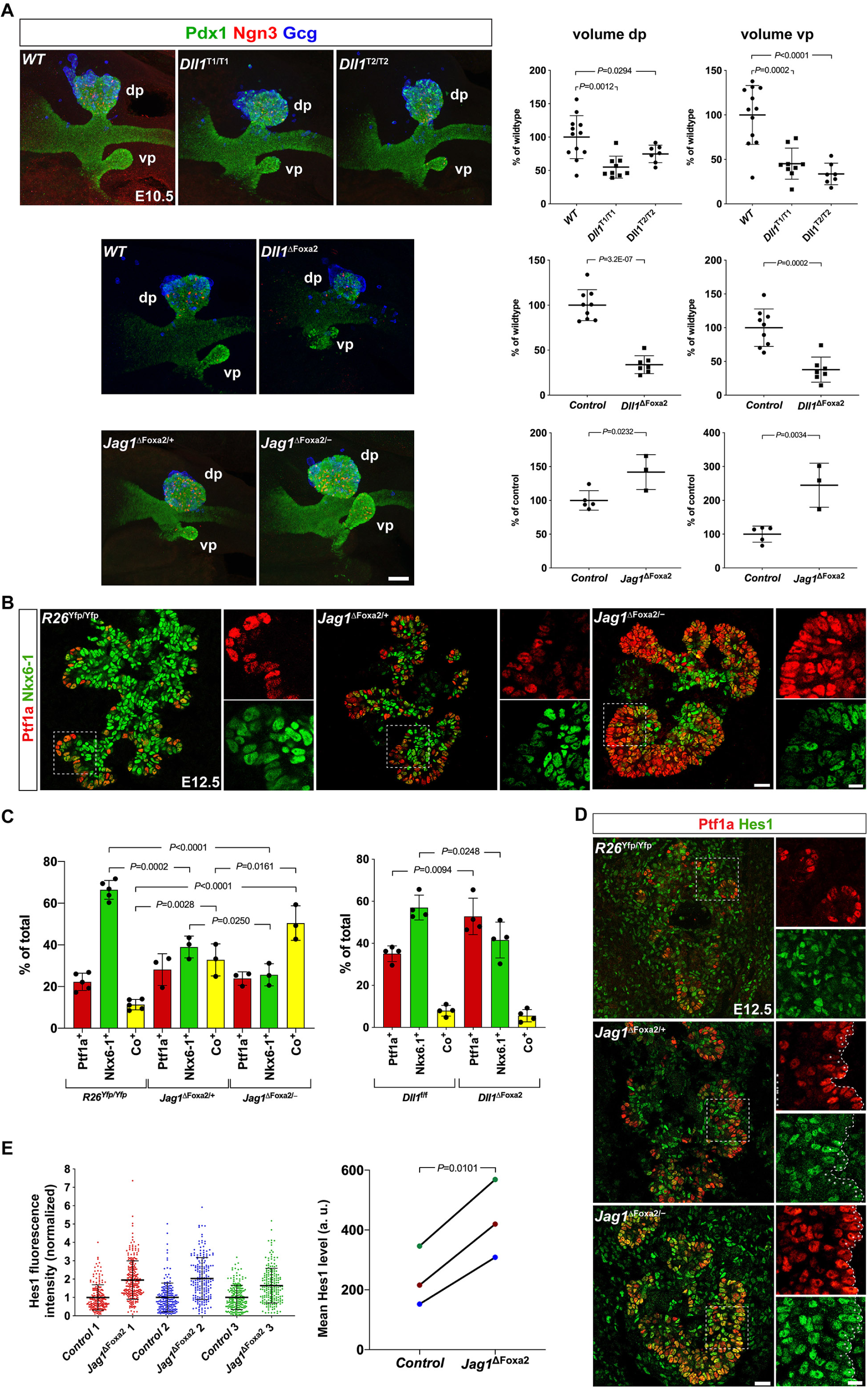
Dll1 and Jag1 differentially affects MPC expansion and differentiation. (A) 3D maximum intensity projections and bud volume quantifications of E10.5 *Dll1*^T1/T1^, *Dll1*^T2/T2^, *Dll1*^ΔFoxa2^, *Jag1*^ΔFoxa2/−^ mutants and their wildtype or heterozygote littermate controls stained by whole-mount IF for Pdx1, Ngn3, and Gcg as indicated. dp: dorsal pancreas; vp: ventral pancreas. Scale bar is 50 µm. (B) Sections of E12.5 wildtype *R26*^Yfp/Yfp^, *Jag1*^ΔFoxa2/+^ and *Jag1*^ΔFoxa2/−^ mutants stained by IF for Ptf1a and Nkx6-1 as indicated. Insets shows higher magnification views of the boxed areas in the main panels. Scale bars, 25 μm (main panels) and 10 μm (insets). (C) Quantification of Ptf1a^+^, Nkx6-1^+^ and Ptf1a^+^Nkx6-1^+^ co-expressing cells in E12.5 wildtype *R26*^Yfp/Yfp^, *Jag1*^ΔFoxa2/+^ and *Jag1*^ΔFoxa2/−^ mutants as well as in Dll1^f/f^ wildtypes and *Dll1*^ΔFoxa2^ mutants. Mean ± S.D., N = 3-5 as indicated by individual data points. (D) Sections of E12.5 *R26*^Yfp/Yfp^, *Jag1*^ΔFoxa2/+^ and *Jag1*^ΔFoxa2/−^ pancreata stained for Ptf1a and Hes1 as indicated. Scale bars are 25 μm (main panels) and 10 μm (insets). (E) Quantification of Hes1 fluorescence intensity in distal Ptf1a^+^ cells in *Jag1*^ΔFoxa2/−^ mutants and *Jag1*^ΔFoxa2/+^ littermate controls. Scatter plots show intensity values for individual cells in three mutant-control pairs in three, separate IF experiments, mean ± S.D., N ≥ 192 cells/embryo. Changes in mean Hes1 levels for each pair of embryos are shown in the right-hand panel, color-coded for embryo ID (a.u., arbitrary units).

We then asked whether bud sizes were affected in E10.5 *Jag1*^ΔFoxa2^ embryos. Since *Foxa2* is linked to *Jag1* on mouse chromosome 2 at a distance of ~5.7 cM we first introduced a *Jag1* null allele (Xue et al., 1999) on the same chromosome as the *Foxa2*^iCre^ allele via meiotic crossover. Animals carrying *Jag1*^−^; *Foxa2*^iCre^ chromosomes were then backcrossed to homozygous *Foxa2*^iCre/iCre^ animals to secure this chromosome from further crossover events. Timed matings of these mice with *Jag1*^fl/fl^; *R26*^YFP/YFP^ animals generated *Jag1*^ΔFoxa2/−^ embryos (referred to as *Jag1*^ΔFoxa2^) and *Jag1*^ΔFoxa2/+^ heterozygote littermate controls. Remarkably, WM-IF analysis revealed the E10.5 dorsal and ventral pancreata to be increased in size in *Jag1*^ΔFoxa2^ embryos compared to controls (Figure 3A). Together, these data suggest that Notch activation is mediated by oscillating Dll1 expression, which via induction of oscillating Hes1 expression stimulates MPC expansion. Conversely, Jag1 attenuates Dll1-mediated Notch activation to limit MPC expansion, possibly by sequestering a fraction of the available Notch receptors in *cis*.

### Suppressing Notch before ~E13 shunts progenitors to a PAC fate

Before testing the requirement for Jag1 and Dll1 in the segregation of MPCs into BP and PAC fates we decided to define the temporal window through which Notch signaling affected this fate choice. To address this question, we administered tamoxifen (Tam) to pregnant dams carrying either *Hnf1b-*CreER^T2^; *Rosa26*^LSL-dnMaml1-eGFP/+^ (hereafter referred to as *R26*^dnMaml1^) embryos or *Hnf1b-*CreER^T2^; *Rosa26*^LSL-YFP/+^ (hereafter referred to as *R26*^YFP^) control embryos at different timepoints and harvested embryos for analysis at E15.5 (Figure 4A). In agreement with a previous *Hnf1b*-CreER^T2^ lineage-tracing analysis (Solar et al., 2009), IF examination of *R26^YFP^* pancreata, treated with Tam between E11.5 – E13.5, for Sox9 and Ptf1a revealed that while some *Hnf1b*-expressing cells retain their multipotency at E13 and E14, they become progressively more biased towards the BP lineage with time (Figures 4B-4D and 4N). In contrast, *R26*^dnMaml1^ pancreata treated with Tam at E11.5 showed an ~11-fold reduction of EGFP^+^ cells adopting a BP fate, while the fraction of EGFP^+^ cells allocated to a PAC fate had increased almost 3-fold. The fraction of labeled cells expressing neither marker was unchanged (Figures 4E and 4N). Similarly, significantly fewer EGFP^+^ cells co-expressed Sox9 in *R26*^dnMaml1^ pancreata injected with Tam at E12.5 or E13.5 compared to controls. However, in sharp contrast to E11.5 Tam-induced pancreata, the proportion of EGFP^+^ cells expressing Ptf1a was unchanged following Tam treatment at E12.5 and E13.5. Instead, more EGFP^+^ cells were Sox9^−^Ptf1a^−^, compared to YFP^+^ cells in the controls (Figures 4F, 4G, 4N). Together, these data show that suppression of Notch signal reception in MPCs and BPs before ~E13 shunts the cells to a PAC fate. However, if Notch signaling is blocked in BPs after ~E13, they adopt an alternative fate.

**Figure 4.**
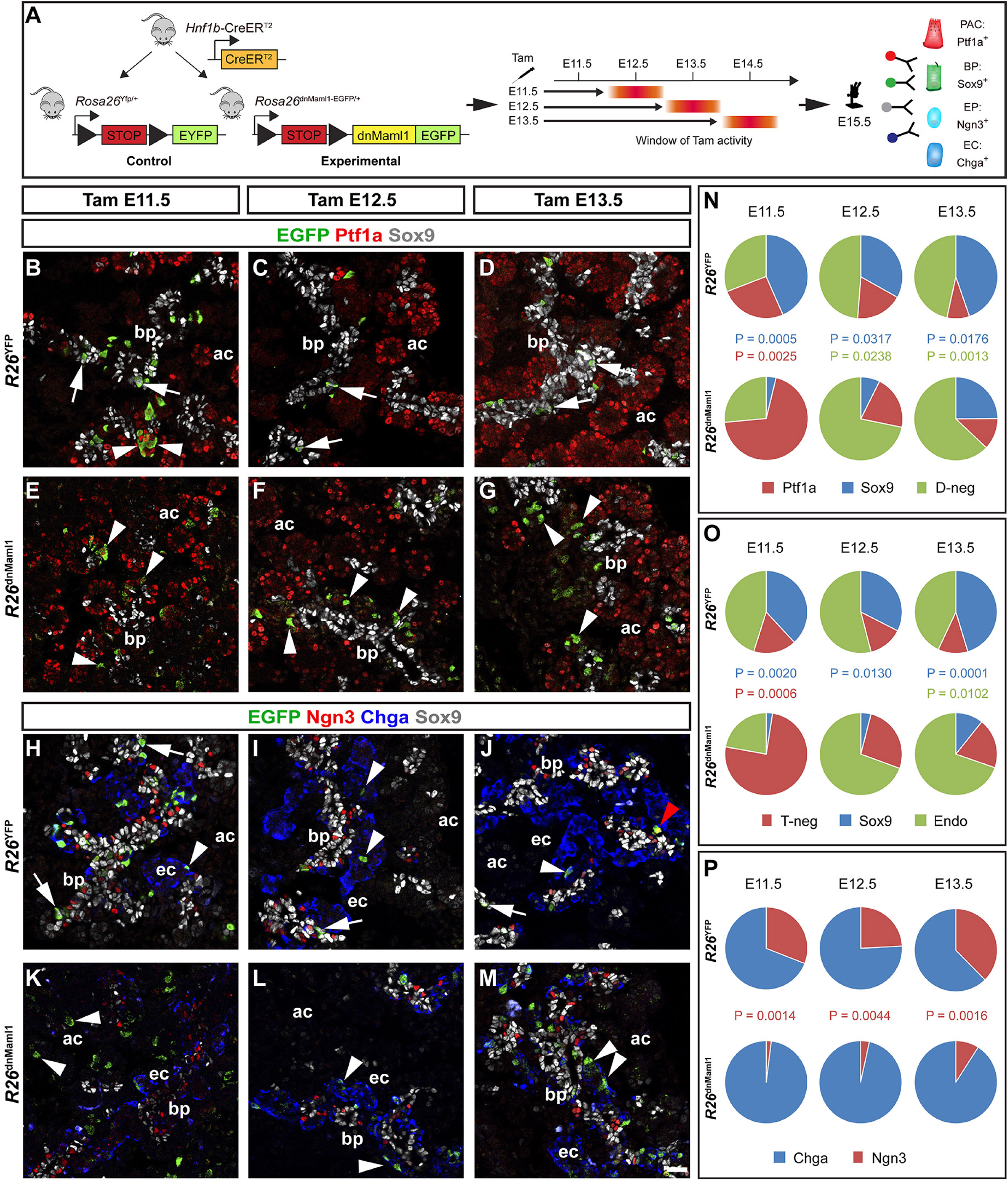
Stage-dependent allocation of progenitor fate by Notch suppression. (A) Overview of strategy applied to identify fates of progeny from pancreatic progenitors. Approximate temporal windows of Tamoxifen (Tam) activity resulting from single intraperitoneal injections at E11.5, E12.5 or E13.5 and cell fate-specific markers are indicated. PAC: pre-acinar cell; BP: bipotent progenitor; EP: endocrine precursor; EC: endocrine cell. (B-M) Sections of E15.5 *Hnf1b-CreER^T2^*; *R26*^YFP^ control (B-D, H-J) or *Hnf1b-CreER^T2^*; *R26*^dnMaml1^ Notch-blocked (E-G, K-M) pancreata stained for EYFP/EGFP and Sox9 combined with either Ptf1a (B-G) or Ngn3 and Chga (H-M), as indicated. bp: bi-potent progenitor domain; ac: emerging acini; ec: endocrine cells. Scale bar 50 μm. Arrows in (B-D, H-J) lineage-labeled Sox9^+^ BPs. Arrowheads in (B, E): lineage-labeled Ptf1a^+^ PACs. Arrowheads in (F, G): Ptf1a/Sox9 double negative cells. Arrowheads in (I-J, L-M) Chga^+^ endocrine cells. Red arrowhead in (J): Ngn3^+^ endocrine precursor. Arrowheads in (K): lineage-labeled Ngn3/Chga/Sox9 triple negative cells. Scale bar is 50 µm. (N-O) Quantification of *R26*^YFP^-(EYFP) *versus R26*^dnMaml1^ (EGFP)-labeled cell distribution at E15.5 following Tam administration at E11.5, E12.5 or E13.5 as indicated. Pie charts show fractions of EYFP/EGFP-labeled cells expressing Ptf1a, Sox9 or neither marker (D-neg) or Sox9, endocrine markers (Endo, Ngn3+Chga) or none of the three (T-neg). (P) Pie chart showing the proportions of labeled cells in the endocrine lineage expressing either Ngn3 or Chga. P-values for significant changes in allocation to distinct cell fates between *R26*^dnMaml1^ and controls are indicated by font color. N = 3 embryos per genotype for each Tam injection time-point. See also Figure S2.

### Suppressing Notch after ~E13 shunts progenitors to an endocrine fate

We next performed IF for EGFP/YFP, Sox9, Ngn3 and Chga to determine when progenitors adopted an endocrine fate upon Notch blockade. As expected, the fraction of labelled cells expressing Sox9 in E15.5 *R26*^dnMaml1^ pancreata was significantly lower than in controls after Tam injection at E11.5 and consistent with the increase of labelled Ptf1a^+^ cells mentioned above, significantly more labelled cells were triple-negative for Sox9, Ngn3 and Chga in dnMaml1 embryos, compared to controls (Figures 4H, 4K, 4O). Examining endocrine lineage markers, we found that the fraction of labelled cells expressing either Ngn3 or Chga, was not significantly different between dnMaml1 embryos and controls (Figures 4H, 4K, 4O). However, fewer labelled cells co-expressed Ngn3 in dnMaml1 embryos than in controls (Figure 4P). Conversely, when Tam was injected at E12.5 and E13.5, the fraction of labelled cells expressing endocrine markers was now markedly increased compared to controls, while the fraction of labelled cells expressing Sox9 was reduced and labelled cells triple-negative for Sox9, Ngn3 and Chga was unchanged between groups (Figures 4I, 4J, 4L, 4M). Again, fewer labelled cells co-expressed Ngn3 in dnMaml1 embryos than in controls (Figure 4P). A qualitative analysis of lineage-traced E15.5 *Sox9*-CreER^T2^; *R26*^dnMaml1^ embryos compared to stage-matched *Sox9-*CreER^T2^; *R26*^YFP^ embryos revealed the same shift in the fate of the labelled cells when comparing Tam injections at E11.5 to E12.5 and E13.5 (Figure S2). Taken together, these results show that the time window through which prevention of Notch activation can shunt cells to a PAC fate closes by ~E13 and confirms that prevention of Notch activation in BPs after ~E13 induces endocrine differentiation (Magenheim et al., 2011; Shih et al., 2012).

### Jag1 is required to exit multipotency

Having established the temporal limit for BP versus PAC specification we next sought to identify the ligands responsible for this fate choice. IF analysis of E12.5 *Jag1*^ΔFoxa2^ mutant pancreas revealed that the fraction of Ptf1a^+^Nkx6-1^+^ MPCs was significantly increased at the expense of Ptf1a^−^Nkx6-1^+^ BPs, while the fraction of Ptf1a^+^Nkx6-1^−^ PACs was unchanged compared to controls (Figures 3B-3C). While Ptf1a is essentially confined to the peripheral-most epithelial cells in wild-type controls, Ptf1a expression was also seen in a few, more proximal cells in *Jag1*^ΔFoxa2/+^ heterozygotes and prominently in many proximal cells in *Jag1*^ΔFoxa2/−^ homozygote mutants (Figure 3B). We previously performed the same analysis on E12.5 *Dll1*^ΔFoxa2^ mutant pancreas (Horn et al., 2012) so we calculated the percentage of each cell type from those data and found that while the fraction of Ptf1a^−^Nkx6-1^+^ BPs was also reduced in these embryos, the fraction of Ptf1a^+^Nkx6-1^+^ MPCs was not increased. Instead, the fraction of Ptf1a^+^Nkx6-1^−^ PACs was increased (Fig. 3C). Thus, proper segregation of MPCs into PAC and BP domains at E12.5 requires Jag1, while Dll1 is required for biasing MPCs towards a BP fate.

Analysis of later stages revealed that most Ptf1a^+^ cells remained co-positive for Nkx6-1 and Sox9 at E13.5, thus maintaining an MPC marker profile. Not until E14.5 did we observe a resolution into distinct Ptf1a^+^Nkx6-1^−^ PACs and Ptf1a^−^Nkx6-1^+^ BPs in the *Jag1*^ΔFoxa2^ pancreas (Figure S3). However, while BPs normally extend all the way into the forming acini at this stage, the *Jag1*^ΔFoxa2^ pancreas was nearly devoid of such cells in the periphery (Figure S3). To begin to understand the mechanism causing MPCs to maintain their multipotent state we assessed how loss of Jag1 impacts Notch activity in *Jag1*^ΔFoxa2^ pancreata by IF analysis of Hes1 expression. We found that Hes1 expression, and by inference active Notch signaling, was upregulated in the E12.5 *Jag1*^ΔFoxa2^ pancreas compared to heterozygote littermates (Figure 3D-3E). This was especially notable in the distal-most cells, in which average Hes1 levels were increased ~2-fold (Figure 3E). This finding suggests that Jag1 is required cell-autonomously to inhibit Notch activation in emerging PACs, and thus act as a symmetry breaker that triggers exit from the multipotent state.

### The combined activities of *Jag1* and *Dll1* specify the entire BP population

Since E15.5 *Mib1*^ΔFoxa2^ mutants are comprised exclusively of PACs cells (Horn et al., 2012) we next examined the individual and combined requirement of Jag1 and Dll1 for BP versus PAC fate at this stage. To address this question, we generated stage-matched *Dll1*^ΔFoxa2^ and *Jag1*^ΔFoxa2^ mutants, compound *Jag1*; *Dll1*^ΔFoxa2^ mutants, and control mice without ligand deletions (*Foxa2*^T2AiCre^*; R26*^YFP^). Analysis of all four genotypes at E15.5 revealed no change in the numbers of PACs or BPs per unit area between single *Dll1*^ΔFoxa2^ mutant and control pancreata. However, the *Dll1*^ΔFoxa2^ mutant pancreata were clearly hypomorphic. In contrast, BPs were severely depleted in *Jag1*^ΔFoxa2^ mutants with a corresponding expansion of PACs, and in compound *Jag1*; *Dll1*^ΔFoxa2^ mutants this shift in fate was essentially all-encompassing (Figures 5A-5E). Notably, Jag1^ΔFoxa2/+^ heterozygotes showed an intermediate phenotype (Figures S4A-S4D). The few remaining BPs found in *Jag1*^ΔFoxa2^ mutants were confined to the central core of the organ (Figure 5A-5B), which was also evident when plotting the fraction of Nkx6-1^+^ and Ptf1a^+^ cells in the *z*-dimension (Figure 5C). IF staining for insulin revealed a ~50% decrease in the number of insulin^+^ β-cells in both *Dll1*^ΔFoxa2^ and *Jag1*^ΔFoxa2^ pancreata compared to controls, while compound *Jag1*; *Dll1*^ΔFoxa2^ mutants exhibited a near complete loss of β-cells (Figure 5D-5E). Together, these data show that Jag1 is required for development of all but the central-most BPs and that the combined activities of Jag1 and Dll1 specify the entire BP lineage and suggest that MPCs normally fated to become BPs adopt a PAC fate in their absence.

**Figure 5.**
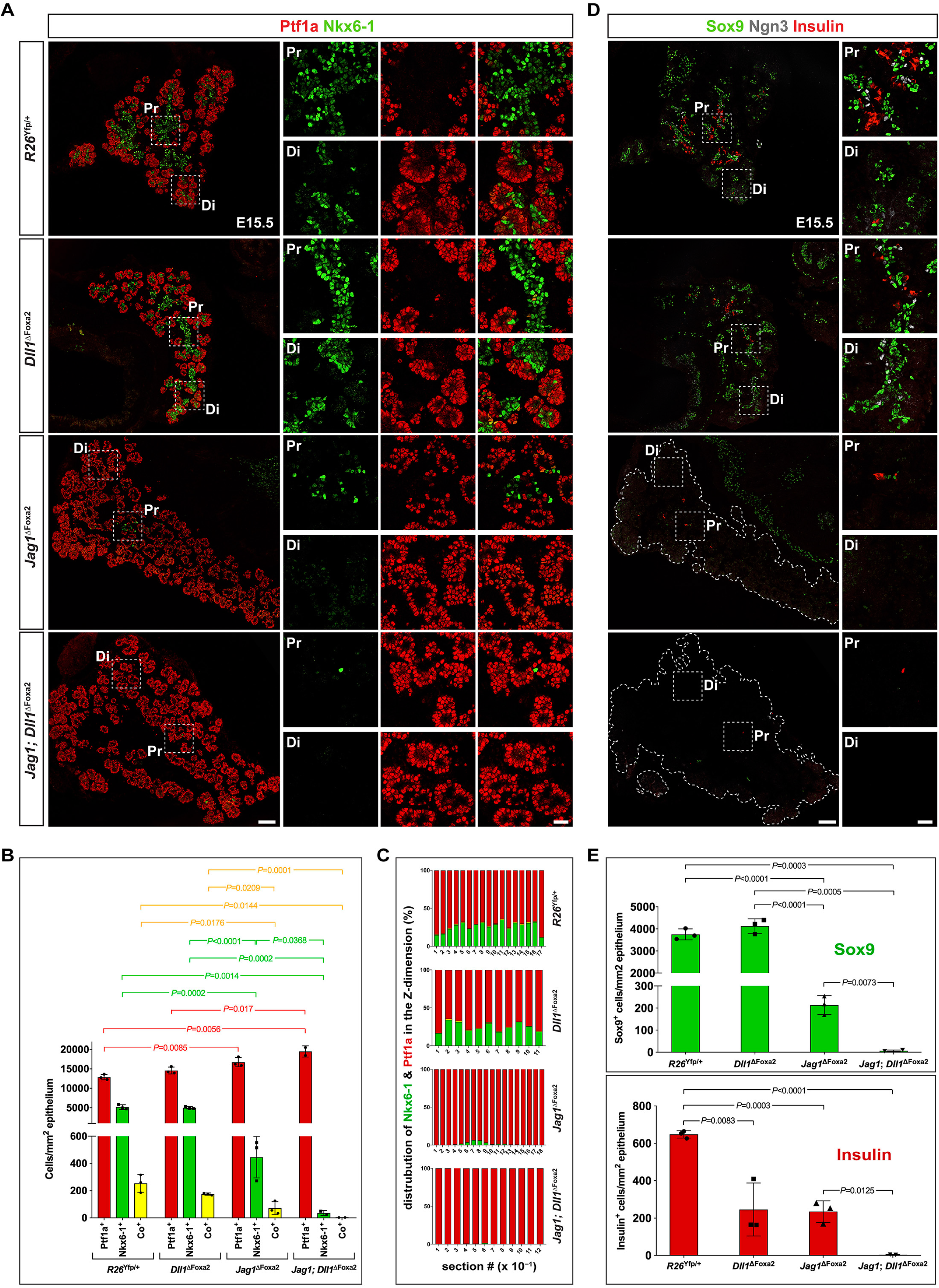
Progenitors adopt a PAC fate in *Jag1-Dll1* double mutant mice. (A) IF for Ptf1a and Nkx6-1 as indicated on sections of E15.5 *R26*^Yfp/+^, *Dll1*^ΔFoxa2^, *Jag1*^ΔFoxa2^, and *Jag1*; *Dll1*^ΔFoxa2^ dorsal pancreata. Di: distal; Pr: proximal. Scale bars are 100 μm (main panels) and 25 μm (insets). (B) Quantitative analyses of the ratio of Ptf1a^+^, Nkx6-1^+^ or Ptf1a^+^Nkx6-1^+^ (Co^+^) cells in E15.5 mutant and control pancreata. (C) Distribution of Ptf1a^+^, Nkx6-1^+^ or Ptf1a^+^Nkx6-1^+^ cells on every 10^th^ section in the *z*-dimension is shown for a representative embryo from each genotype. (D) IF for Sox9, Ngn3 and insulin as indicated on sections of E15.5 *R26*^Yfp/+^, *Dll1*^ΔFoxa2^, *Jag1*^ΔFoxa2^, and *Jag1*; *Dll1*^ΔFoxa2^ dorsal pancreata. Di: distal; Pr: proximal. Scale bars are 100 μm (main panels) and 25 μm (insets). (E) Quantitative analysis of Sox9^+^ and insulin^+^ β-cells in E15.5 mutant and control pancreata. Graphs show mean ± S.D., N = 3 except for *Jag1*; *Dll1*^ΔFoxa2^ where N = 2. See also Figures S3, S4 and S5.

### Impeding ligand trans-activation in late development only affects CACs

The strong phenotype we observe in *Jag1*; *Dll1*^ΔFoxa2^ mutant embryos contrasts with the rather modest phenotype seen in *Jag1*; *Dll1*^ΔPtf1a^ mutant embryos, which only lack CACs (Nakano et al., 2015). This suggests that the timing of Cre-mediated recombination, which is mosaic and occurs considerably later with *Ptf1a*^Cre^ compared with *Foxa2*^iCre44,48^, is critical for deciding the outcome. Considering that Mib1 is required for all Notch ligand trans-activation (Itoh et al., 2003; Koo et al., 2005; Koo et al., 2007) and that its elimination in *Mib1*^ΔFoxa2^ pancreata phenocopies the cell fate changes in *Jag1*; *Dll1*^ΔFoxa2^ embryos seen here (Horn et al., 2012), we therefore asked whether *Mib1*^ΔPtf1a^ pancreata would phenocopy *Jag1*; *Dll1*^ΔPtf1a^ pancreata. *Mib1*-depleted cells and their progeny were identified by *R26*^YFP^ recombination. Examination of *Mib1*^ΔPtf1a^ pancreata prior to E13.5 failed to reveal any obvious defects. In contrast, analysis at E15.5 revealed the ductal tree to be truncated distally. The distal-most, Sox9^+^ cells (prospective CACs), which normally protrude into the nascent Ptf1a^+^ acini are specifically depleted in *Mib1*^ΔPtf1a^ embryos (Figure S5). Notably, this phenotype closely resembles the reported loss of CACs following *Ptf1a*^Cre^-driven compound deletion of *Dll1* and *Jag1* (Nakano et al., 2015). Taken together, these results suggest that the trans-activation of Notch receptors by Jag1 and Dll1 expressed on PACs and emerging acinar cells is required to specify and/or maintain adjacent CAC precursors.

### Coupling of Hes1 oscillation parameters to cell fate

Similar to *Jag1*; *Dll1*^ΔFoxa2^ mutants, *Hes1*^ΔFoxa2^ embryos also show severely reduced numbers of BPs and a corresponding increase in PACs (Horn et al., 2012). We therefore asked whether perturbing the period or amplitude of Hes1 oscillations would similarly affect cell fate. To address this question, we exploited the ability of NICD levels to modulate oscillation parameters (Wiedermann et al., 2015). We explanted E10.5 dorsal buds from *Hes1*-Luc2 embryos and perturbed NICD levels with small molecule inhibitors. We monitored the period and amplitude of Hes1 oscillation by bioluminescence imaging and conducted end-point IF analysis of cell fate allocation. Western blot analysis showed that treatment with a low dose of the γ-secretase inhibitor DAPT reduced average N1ICD levels by ~50% compared to vehicle controls (Figure 6A) and bioluminescence imaging revealed a ~70% decrease in the amplitude of Hes1 oscillations, while the period was unaffected (Figure 6B). This was associated with an increase of Ptf1a^+^ PACs and Ptf1a^+^Sox9^+^ MPCs at the expense of Sox9^+^ BPs at the end of the 4-day culture period (Figure 6C-6D). In contrast, exposure to MLN4924, a Nedd8 activating enzyme inhibitor, increased N1ICD levels by ~60% compared to vehicle controls (Figure 6A) and prolonged the mean oscillation period from ~90 to ~120 minutes while only marginally affecting the amplitude (Figure 6B). IF analysis showed that Ptf1a^+^ cells (PACs and MPCs) were strongly reduced by MLN4924 treatment and the explants were composed almost entirely of Sox9^+^ cells (BPs and/or duct cells) (Figure 6C-6D). Together these data suggest that the both the amplitude and period of Hes1 oscillations are important for MPC fate choice.

**Figure 6.**
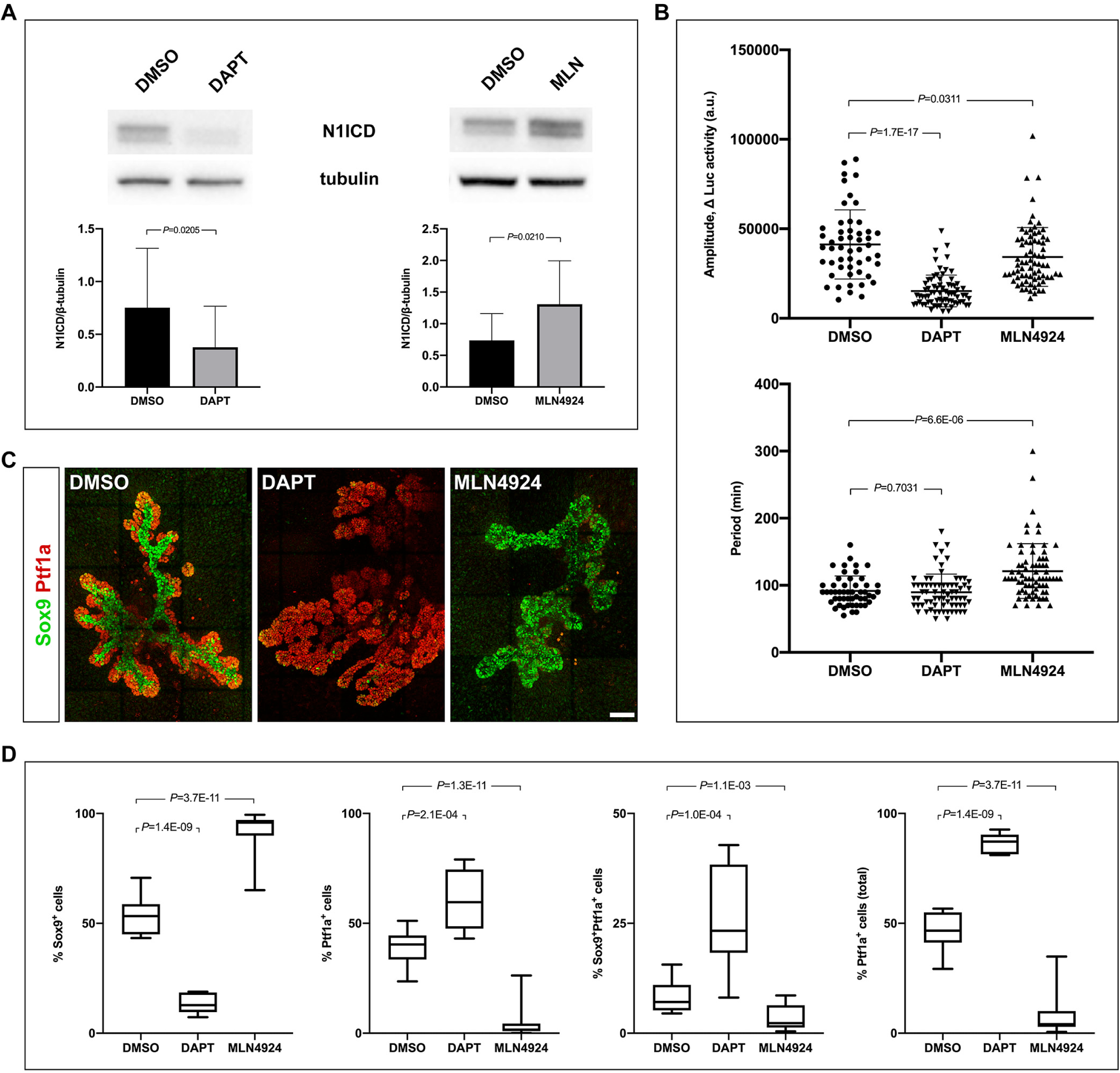
Modulation of NICD levels affect Hes1 oscillation parameters and cell fate. (A) Western blots showing reduced or elevated N1ICD levels in response to DAPT (2.5 µM) or MLN4924 (5 µM) treatment, respectively. Bar graph shows quantification of band intensities from individual experiments (mean ± S.D., N = 4). (B) Distributions and averages of the of amplitudes and periods for Hes1 oscillations in E10.5 pancreas explants treated with DAPT or MLN4924. (Mean ± S.D., N > 50 cells for each condition. (C) 3D maximum intensity projections of E10.5 pancreas explants cultured for 5 days in DAPT or MLN4924 and stained by whole-mount IF for Sox9 and Ptf1a. Scale bar is 100 µm. (D) Quantitative analyses of the ratio of Ptf1a^+^, Sox9^+^ or Ptf1a^+^Sox9^+^ cells from E10.5 pancreas explants cultured as in (C). Median and interquartile ranges as well as minimum and maximum values are indicated, N = 13 for DMSO- and MLN4924-treated samples and N = 7 for DAPT treated samples. See also Figure S6.

### Early *Jag1*^ΔFoxa2^ mutants have normal plexus formation and organ architecture

We next analyzed how the BP-to-PAC fate switch affects ductal morphogenesis and overall organ development in *Jag1*^ΔFoxa2^ mutants. In spite of the delayed PD patterning, we found that formation of the epithelial plexus occurred normally and overall organ size and morphology was comparable between *Jag1*^ΔFoxa2^ mutants and heterozygote littermate controls (Figure 7A). Remarkably, even at E15.5 the overall organ architecture is essentially unaffected. The dorsal pancreas had a well-formed, anvil-shaped head and the body gradually tapered into the narrow connection with a normal-sized ventral pancreas located in the duodenal loop (Figure 7B). In contrast, the *Dll1*^ΔFoxa2^ dorsal pancreas was hypoplastic with the head being malformed and the body of the pancreas being greatly truncated (Figure 7B). However, closer inspection of *Jag1*^ΔFoxa2^ embryos revealed that although the ductal plexus appeared to have remodeled into a hierarchical tree-like structure, the smooth walls of the intercalated ducts seen in controls and *Dll1*^ΔFoxa2^ mutants, showed a more serrated appearance in *Jag1*^ΔFoxa2^ mutants (Figure 7B). Nevertheless, acinar structure appears normal with apical localization of Muc1, ZO-1 and PKCζ, indicating that acinar cytoarchitecture is maintained in the E15.5 *Jag1*^ΔFoxa2^ pancreas (Figure 7H-7I). In the domain usually occupied by Sox9^+^ prospective ducts, we were able to identify elongated tubular structures expressing Ptf1a instead of Sox9 (Figure S4E). Similar to normal PACs these mis-specified PACs also expressed Bhlha15/Mist1 (Figure S4F). Taken together, these findings suggest that the overall organ architecture and remodeling of the ductal plexus is regulated independently from the differentiation programs that allocate MPCs to endocrine, duct and acinar lineages. In contrast, finer morphological features of the ductal tree are clearly perturbed by the BP-to-PAC fate switch.

**Figure 7.**
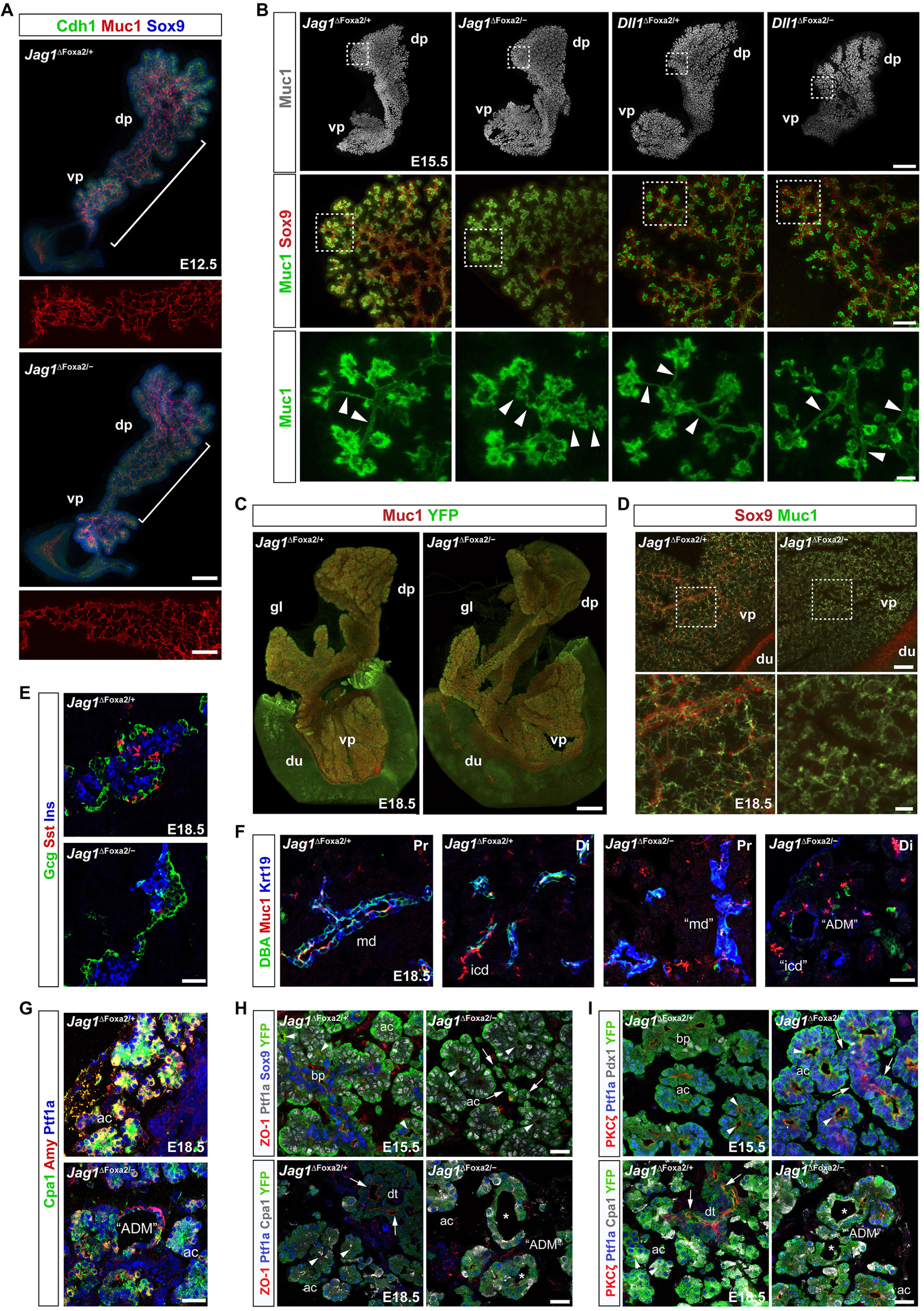
Ductal remodeling is defective in *Jag1*^ΔFoxa2^ mutants. (A) 3D maximum intensity projections of the midgut region of E12.5 *Jag1*^ΔFoxa2/+^ and *Jag1*^ΔFoxa2/−^ littermates stained for Cdh1, Muc1 and Sox9 as indicated by whole-mount IF. White brackets indicate the Muc1 staining in the body region of the dorsal pancreata to emphasize the luminal plexus regions shown in insets below the main panels. dp: dorsal pancreas, vp: ventral pancreas. Scale bars are 50 µm (main panels) and 25 µm (insets). (B) Upper panels: 3D maximum intensity projections of whole-mount, Muc1 labeled E15.5 *Jag1*^ΔFoxa2/+^, *Jag1*^ΔFoxa2/−^, *Dll1*^ΔFoxa2/+^, and *Dll1*^ΔFoxa2/−^ pancreata. Scale bar is 300 µm. Middle panels: Higher power images of maximum intensity projections from sub-stacks acquired from the boxed areas indicated in upper panels showing Muc1 and Sox9 distribution. Scale bar is 50 µm. Lower panels: Higher power images of boxed areas indicated in middle panels showing Muc1 alone to emphasize the structure of the terminal ducts. Arrowheads: Terminal duct lumens appear serrated in *Jag1*^ΔFoxa2/−^ mutants while heterozygote littermate controls and *Dll1*^ΔFoxa2/−^ mutants show smooth lumens. dp: dorsal pancreas, vp: ventral pancreas. Scale bar is 10 µm. (C) 3D maximum intensity projections of E18.5 *Jag1*^ΔFoxa2/+^ and *Jag1*^ΔFoxa2/−^ whole-mount pancreata stained for Muc1 and YFP as indicated. dp: dorsal pancreas, gl: gastric lobe, vp: ventral pancreas, du:duodenum. Scale bar is 700 µm. (D) 3D maximum intensity projections of the same pancreata shown in (C). Note absence of both interlobular and large intralobular ducts in *Jag1*^ΔFoxa2/−^ mutants compared to controls. Boxed areas indicate regions shown at higher magnifications in lower panels. Scale bars are 100 µm (upper panels) and 25 µm (lower panels). (E-G) Sections of E18.5 *Jag1*^ΔFoxa2/+^ and *Jag1*^ΔFoxa2/−^ pancreata stained for insulin (Ins), glucagon (Gcg) and somatostatin (Sst) (E), Muc1, Krt19 and DBA (F), or Cpa1, amylase (Amy) and Ptf1a as indicated (G). md: main duct, (disrupted in mutants), icd: intercalated ducts (abnormal morphology in mutants), “ADM”: ADM-like structures. Scale bars are 25 µm. (H-I) Sections of E15.5 and E18.5 *Jag1*^ΔFoxa2/+^ and *Jag1*^ΔFoxa2/−^ pancreata stained for ZO-1, Ptf1a, Sox9 and YFP or ZO-1, Ptf1a, Cpa1 and YFP as indicated (H) and PKCζ, Ptf1a, Pdx1 and YFP or PKCζ, Ptf1a, Cpa1 and YFP as indicated (I). Arrows: Ptf1a^+^ duct-like structures in E15.5 *Jag1*^ΔFoxa2/−^ mutants and apical ZO-1 and PKCζ in E18.5 duct cells in control mice. Arrowheads: apical ZO-1 in the E15-5 Ptf1a^+^ PAC domain and in E18.5 Ptf1a^+^Cpa1^+^ acinar cells. Asterisks: Absence of apical ZO-1 and PKCζ in YFP^+^Ptf1a^+^Cpa1^+^ ADM-like structures (“ADM”) in E18.5 *Jag1*^ΔFoxa2/−^ pancreas. Scale bars, 25 µm. ac: acini, bp: bi-potent progenitor domain, dt: ducts.

### Late *Jag1*^ΔFoxa2^ embryos show signs of acute pancreatitis and ADM

E18.5 *Jag1*^ΔFoxa2^ pancreata remained equivalent to controls in size and overall organ morphology. The dorsal and ventral pancreas appeared fused normally, the gastric lobe was present, and the normal “anvil” shape of the dorsal pancreas (Villasenor et al., 2010) was evident (Figure 7C). However, closer inspection revealed that the larger ducts were not formed properly. The main duct was disrupted, interlobular as well as larger intralobular ducts were largely absent and many of the terminal ducts appeared serrated, occasionally connected by ducts of relatively normal morphology despite being composed of Ptf1a^+^ cells (Figure 7D). As expected, we saw a prominent loss of endocrine cells in E18.5 *Jag1*^ΔFoxa2^ pancreata, with scattered α-cells and a central cluster of β-cells. The late-arising, somatostatin^+^ δ-cells were nearly absent, even in the central cluster (Figure 7E). Examination of Muc1 and Krt19 expression and DBA lectin binding revealed an obvious paucity of the ductal tree and throughout the epithelium we observed numerous “ring-like” structures with perturbed apicobasal polarity and co-expression of amylase, Cpa1 and Krt19 (Figure 7F-7I), reminiscent of the acinar-to-ductal metaplasia (ADM) associated with Cerulein-induced acute pancreatitis (Nishikawa et al., 2019).

## Discussion

Regulation of MPC proliferation and fate choice by Notch has been comprehensively documented by previous work (Afelik et al., 2012; Ahnfelt-Rønne et al., 2012; Apelqvist et al., 1999; Fujikura et al., 2006; Fujikura et al., 2007; Hald et al., 2003; Jensen et al., 2000a; Jensen et al., 2000b; Murtaugh et al., 2003; Schaffer et al., 2010; Shih et al., 2012). In this study we show that the Notch pathway components Dll1 and Hes1 display ultradian oscillations in the developing mouse pancreas and that Jag1, which is uniformly expressed in MPCs, dampens Dll1-mediated Notch activation to restrain MPC growth, secure timely exit from the multipotent state, and coordinate MPC fate choice (Figure S6).

We found the period of pancreatic Dll1 and Hes1 oscillations to be in the range of 1-2 hours with an average of ~90 minutes. This is noticeably shorter than the period observed in neural progenitors, which is in the range of 2-3 hours with an average of ~150 minutes (Imayoshi et al., 2013; Shimojo et al., 2016). At first glance this may appear surprising given that the regulatory network between and within cells has a similar architecture for neural and pancreatic progenitors (this work and (Kageyama et al., 2008; Tiedemann et al., 2017)). Moreover, the oscillations in neighboring pancreatic and neural progenitors are typically not in phase, while groups of oscillating cells in the pre-somitic mesoderm *are* in phase. Indeed, we often observed Hes1 oscillations in neighboring pancreatic cells to be in anti-phase, as is also observed in neural progenitors (Imayoshi et al., 2013). So how can the different periods be explained? The oscillatory period in pre-somitic mesoderm is sensitive to Notch activity and modeling suggests that this may be a general property of Notch oscillations (Kim et al., 2011; Wiedermann et al., 2015). We therefore speculate that the high expression levels of Notch pathway components in neural progenitors compared to MPCs may cause the former to attain a longer period. This notion is supported by our observation of an extended period in pancreatic progenitors treated with MLN4924 but raises the question of why the period is not shortened by DAPT treatment. However, a minimal period is imposed on the system, partly by the sum of the transcriptional and translational delays in the delayed negative feedback mechanism of Hes1 and partly by the parameters governing the dynamics of Dll1/Notch-mediated intercellular signaling (Lewis, 2003), and it is possible that a minimal period is reached in the pancreas.

We have previously shown that E10.5 *Dll1*^−/−^ pancreata are hypoplastic due to reduced proliferation (Ahnfelt-Rønne et al., 2012) and now we show that E10.5 *Dll1*^ΔFoxa2^ mutants and *Dll1* Type 1 and Type 2 oscillation mutants also present with pancreatic hypoplasia. Both Type 1 and Type 2 mutants encode the wildtype protein, but due to accelerated or delayed protein synthesis, respectively, a steady intermediate level of Dll1 expression is achieved as oscillations of both Dll1 and Hes1 are dampened (Shimojo and Kageyama, 2016). The hypoplasia seen in these mutants suggests that oscillatory expression of Dll1 and/or Hes1 is important for stimulating MPC growth. This can be explained if ultradian Dll1 oscillations enables a temporal symmetry where MPCs alternate between sending and receiving input via Notch. Without oscillations it would prove difficult for the cells to reach a ligand^Lo^, receiving state as they would be subject to *cis*-inhibition and fail to receive input via Notch receptors that ultimately couples to the mitotic machinery (Figure S6A).

Notably, we found that E10.5 Jag1-deficient pancreata were hyperplastic, showing that Jag1 limits MPC growth. Together, these results suggest that Jag1 antagonizes Dll1 function at this stage, possibly by a *cis*-inhibitory interaction that sequesters a fraction of the available Notch receptors (Figure S6A). It is possible that Jag1-mediated dampening of Notch activation is part of the mechanism that sets the oscillatory frequency of Hes1 and Dll1 expression.

In pancreata undergoing PD patterning, we found that emerging PACs (Ptf1a^+^) expressed high levels of ligands and little to no Hes1, indicating a state of low Notch activity. Conversely, adjacent MPCs (Ptf1a^+^Nkx6-1^+^) or BPs (Nkx6-1^+^) were generally expressing no or low levels of ligand and high levels of Hes1, indicating Notch activation. This suggests that mutually inactivating *cis*-interactions between ligands and receptors (Sprinzak et al., 2010) are crucial for exiting the multipotent stage and for the cells to adopt either a PAC or BP fate. Such a notion is supported by the downregulation of Hes1 and absence of Notch2 expression in PACs and thus overall lower levels of Notch receptor expression than seen in emerging Notch1^+^Notch2^+^ BPs. Conversely, Notch1/2 co-expression, and the absence of Jag1 expression in nascent BPs, would render these more sensitive to signal reception and less prone to signal emission due to *cis* interactions (Figure S6B).

To test these ideas and to investigate the role of individual ligands in PD patterning we performed single and double Dll1/Jag1 loss-of-function experiments. Our marker analyses showed that most epithelial cells in the E12.5 *Jag1*^ΔFoxa2^ embryos maintained a Ptf1a^+^Nkx6-1^+^Sox9^+^ marker profile suggesting that they failed to exit the MPC stage. This correlated with increased Hes1 expression, suggesting that Notch activity is increased and that Jag1 normally acts cell-autonomously to inhibit Notch activation in emerging PACs. We suggest that upregulation of Jag1 *cis*-inhibits Notch receptors and thus acts as a symmetry breaker that terminates oscillatory Hes1 expression in nascent PACs (Figure S6B). Loss of Notch activity may additionally downregulate Nkx6-1 (Afelik et al., 2012) and/or liberate Rbpj from N1ICD, which would then be free to complex with Ptf1a (Cras-Meneur et al., 2009). Both of these mechanisms would favor a PAC fate (Cras-Meneur et al., 2009; Schaffer et al., 2010). Concurrently, free ligand molecules would be able to convey *trans*-activation of receptors in neighboring cells if these are in a responsive, ligand^Lo^/receptor^Hi^ state, and instruct these to adopt a BP fate (Figure S6C).

In spite of increased Notch activity at early stages, *Jag1*^ΔFoxa2^ MPCs eventually adopt a PAC fate, which depends on the cells attaining a state of low Notch activation (Afelik et al., 2012; Horn et al., 2012; Schaffer et al., 2010). However, it remains to be determined what triggers a reduction in Notch activation, but it is noteworthy that the timing coincides with the onset of *Lfng* (*Lunatic fringe*) expression at E14.5 in PACs (Svensson et al., 2009). Lunatic fringe has been shown to inhibit Notch activity in presomitic mesoderm (Dale et al., 2003) and could potentially strengthen *cis*-inhibitory activity of Dll1 (LeBon et al., 2014) in E14.5 *Jag1*^ΔFoxa2^ progenitors. This would attenuate expression of the BP-promoting Notch target genes Nkx6-1 (Afelik et al., 2012) and Sox9 (Shih et al., 2012) in Dll1^Hi^ cells allowing these to exit from the MPC state and ultimately adopt a PAC fate. Testing this hypothesis and identifying the precise role of different Notch receptors awaits future experiments.

The strong effect on PD patterning we observe is surprising since previous analyses of pancreas-specific Jag1 or Jag1/Dll1 deletions did not uncover a prominent expansion of the PAC domain at the expense of BPs (Golson et al., 2009a; Golson et al., 2009b; Nakano et al., 2015). We suspect that this can be attributed to the different timing of efficient, non-mosaic recombination between different Cre lines. Conditional Jag1 deletion with *Pdx1*-Cre or *Foxa3*-Cre driver lines occurs much later than with our targeted *Foxa2*^Cre^ driver, which recombined with >99% efficacy prior to pancreas specification (Horn et al., 2012). More recently, compound *Dll1*; *Jag1*^ΔPtf1a^ mutants were found to have a loss of CACs, the terminal-most cell type in the ductal tree, while single mutant littermates did not show any phenotype (Nakano et al., 2015). As our *Mib1*^ΔPtf1a^ mutants phenocopy the loss of CACs reported in *Dll1*; *Jag1*^ΔPtf1a^ mutants, this suggests that the *Ptf1a*^Cre^-driver only becomes non-mosaic in PACs and their progeny and that CACs are specified by PACs late in pancreatic development.

In spite of the prolonged MPC state, the ductal plexus forms normally. However, at later stages the consequence of the BP-to-PAC switch becomes evident as normal remodeling of the ductal plexus into a well-structured ductal tree is disrupted. The tubular network comprising the intercalated ducts seems to form but the normal smooth morphology of the ductal lumen is perturbed by E15.5. We also noted a complete absence of larger intralobular ducts and interlobular ducts at E18.5 and ductal structures in the area of the main duct appear interrupted as also previously noted in *Jag1*^ΔPdx1^ animals (Golson et al., 2009b). These perturbations are not surprising given that acinar cells have not evolved to form cuboidal or columnar epithelia, but rather to adopt a pyramidal shape fitting for cells forming an acinus. However, in spite of these disturbances the overall organ morphology is surprisingly well preserved. We conclude that the regulatory principles governing the overall shape of the pancreas are highly resilient to cell fate changes, at least as long as these occur in the internal part of the organ.

## Acknowledgements

We thank Ole D. Madsen and Jane E. Johnson for antibodies, Young-Yun Kong, Jorge Ferrer, Heiko Lickert, and Chris V. E. Wright for mouse lines, and Thi Nguyen for technical assistance. P.S. received grants from the Novo Nordisk Foundation (NNF16076 and NNF10717). The Novo Nordisk Foundation Center for Stem Cell Biology is supported by a Novo Nordisk Foundation grant number NNF17CC0027852.

## Author Contributions

P.A.S., C.A.C and A.l.R.E-M designed, carried out experiments and wrote the manuscript. M.C.J. performed all staining, imaging and quantitative analysis of whole-mount specimens. K.H.L. contributed to phenotype analysis. H.S., I.I. and R.Ka. contributed to bioluminescence imaging and generated the *Jag1*^J1VmC^ and *Dll1*^D1VmC^ mouse lines. R.Ko. generated the N1IP::Cre^HI^; *Rosa26*^LSL-Ai3^ embryos. P.S. conceived the study, designed and interpreted experiments and wrote the manuscript. All authors revised and approved the manuscript.

## Methods

### Animals

Published mouse strains were genotyped according to the original work: *R26*^LSL-dnMaml1-EGFP^ (Horn et al., 2012), *Gt(ROSA)26Sortm1(EYFP)Cos* (*R26*^LSL-YFP^ reporter (Srinivas et al., 2001)), *Gt(ROSA)26Sortm3*(CAG-EYFP)Hze (*R26*^LSL-Ai3^ reporter (Madisen et al., 2010)), *Mib1*^tm2Kong^ (floxed *Mib1* (Koo et al., 2007)), *Dll1*^tm1Gos^ (*Dll1*^LacZ^ null allele (Hrabe de Angelis et al., 1997))*, Dll1*^tm1.1Hri^ (floxed *Dll1* (Horn et al., 2012)), *Jag1*^tm1Grid^ (*Jag1* null allele (Xue et al., 1999)), *Jag1*^tm2Grid^ (floxed *Jag1* (Kiernan et al., 2006)), Tg(Hes1-EGFP)^1Hri^ (BAC transgenic *Hes1-EGFP* reporter (Klinck et al., 2011)), *Hes1*^tm1Fgu^ (*Hes1* null allele (Ishibashi et al., 1995)), *Foxa2*^T2AiCre^ (Cre add-on allele (Horn et al., 2012)), *Hnf1b-*CreER^T2^ (Solar et al., 2009), *Sox9-*CreER^T2^ (Kopp et al., 2011), *Ptf1a*^Cre^ (Kawaguchi et al., 2002) and *Notch1*^tm4(cre)Rko^ (N1IP::Cre^HI^ (Liu et al., 2015)), Luc2-Hes1 (BAC transgenic Luciferase-Hes1 fusion protein reporter (Imayoshi et al., 2013)), and lastly *Dll1*^tm1.1Kag^ (*Dll1*-Fluc), *Dll1*^tm4.1(Dll1)Kag^ (*Dll1* Type 1 mutant), and *Dll1*^tm5.1(Dll1)Kag^ (*Dll1* Type 2 mutant) all from (Shimojo and Kageyama, 2016). Additional genotyping primers are given in Supplementary Table 1. Homozygous *Dll1*^D1VmC^ and *Jag1*^J1VmC^ mice are viable and fertile, but were maintained and analyzed as heterozygotes due to *Dll1*^D1VmC^ being a weak hypomorph evident by short, kinky tails in homozygote animals. Generation of Jag1 C-terminal Venus-T2A-mCherry fusion reporter knock-in construct was conducted using a BAC clone (RP23-173O12) from the BACPAC Resources Center at Children’s Hospital Oakland Research Institute. An frt-PGK-EM7-Neo-frt cassette was inserted downstream of a Venus-T2A-mCherry reporter in pBluescript II SK+, flanked by 300-500-bp homology arms from the *Jag1* gene with the *Jag1* stop codon removed. BAC targeting cassettes were excised and electroporated into competent SW105 cells containing the BAC clone of interest. Correctly targeted BAC clones were identified by a panel of PCR primers and restriction digestions. The knock-in cassette fragment was retrieved and cloned into pMCS-DTA (a kind gift from Dr. Kosuke Yusa, Osaka University, Japan). The 5’- and 3’-homology arms in the retrieval vector were designed such that 2.5- and 7.5-kb DNA segments flanked the Venus-T2A-mCherry reporter-frt-PGK-EM7-Neo-frt cassette in the BAC clone, which was then subcloned into pMCS-DTA. The shorter homology arm was used to design PCR-based screening for targeted ES cells (TT2). Chimeric mice were produced from successfully targeted ES cell clones by aggregation with ICR embryos. Germ line transmission of the targeted allele was assessed by PCR of tail DNA. pCAG-FLPe mice (Kanki et al., 2006) were used to remove the frt-PGK-EM7-Neo-frt cassette. The Dll1-Venus-T2A-mCherry knock-in mice were generated by a similar strategy using a *Dll1* containing BAC clone (RP23-306J23).

*Jag1*^ΔFoxa2^ and *Dll1*; *Jag1*^ΔFoxa2^: We first introduced a *Jag1* null allele (Xue et al., 1999) on the chromosome carrying the *Foxa2*^iCre^ allele via meiotic crossover. Animals carrying *Jag1*^−^; *Foxa2*^iCre^ chromosomes were then backcrossed to homozygous *Foxa2*^iCre/iCre^ animals to secure this chromosome from further crossover events. *Jag1*^+/−^; *Foxa2*^iCre/iCre^ animals were next crossed with *Jag1*^fl/fl^*R26*^YFP/YFP^ animals to generate *Jag1*^ΔFoxa2^ embryos and to *Dll1*^fl/+^; *Foxa2*^T2AiCre/T2AiCre^ animals to generate *Jag1*^+/−^*; Dll1*^fl/+^*; Foxa2*^T2AiCre/T2AiCre^ mice. The latter was then crossed with *Jag1*^fl/fl^; *Dll1*^fl/fl^; *R26*^YFP/YFP^ animals to generate *Jag1*; *Dll1*^ΔFoxa2^ embryos. *Dll1*^ΔFoxa2^ embryos and *R26*^YFP^ controls were made as previously described (Horn et al., 2012). Noon on the day of vaginal plug appearance was considered E0.5. All animal experiments described herein were conducted in accordance with local legislation and authorized by the local regulatory authorities.

### Tamoxifen administration

Tamoxifen (Sigma-Aldrich, St. Louis, MO) was dissolved at 10 mg/ml in corn oil (Sigma) and a single dose of 75 μg/g (for *Hnf1b-* and *Sox9-*CreER^T2^-mediated *R26*^YFP/dnMaml1-eGFP^ induction) or 40 μg/g (for *Sox17*^CreERT2^-mediated ligand deletion) body-weight administered by intraperitoneal injection at noon ± 1 hour.

### Explant culture

Dorsal pancreata from luciferase reporter embryos where isolated at E10.5 or E12.5 and cultured on 35 mm glass-bottom µ-dishes (Ibidi) coated with 0.1 mg/mL Fibronectin (Sigma-Aldrich, St. Louis, MO) in explant culture medium: M199 (Gibco?) supplemented with 10% FCS, 1% Pen/strep, 1% Fungizone, 25 ng/ml EGF, 25 ng/ml FGF2 and 100 ng/ml FGF10 (R&D Systems). The explants were incubated at 37°C, 20% O_2_ and 5% CO_2_. The explants dissected at E10.5 were cultured for various durations as indicated in the main text. The Dll1-Luc2 explants dissected at E12.5 were cultured for 4 hours before over-night (20 hours) imaging was initiated.

### Western blots

8-10 dorsal pancreata from E10.5 embryos were pooled and cultured for three hours in hanging drops of explant culture medium supplemented with either 2.5 µM DAPT (Sigma-Aldrich, St. Louis, MO), 5 µM MLN4924 (Cayman Chemical, Ann Arbor, MI) or DMSO. Explants were lysed in lysis buffer (Cell Signaling Technology, Leiden, NL). Samples were sonicated and spun at 15.000 rpm for 20 min at 4°C, then reduced at 95°C for 10 min and loaded on a Bolt^TM^ 4-12% gel Bis-Tris Plus acrylamide gel (Thermo Fischer Scientific, Slangerup, DK). Protein was transferred using a Novex™ Semi-Dry Blotter (InVitrogen, Slangerup, DK) according to the manufacturers instructions and blocked in blocking buffer (5% milk in PBST (PBS w. 0.1% Tween20)). Membranes were incubated overnight at 4°C in primary antibodies diluted in blocking buffer and after washing in PBST the blots were developed with the SuperSignal West Dura ECL kit (InVitrogen, Slangerup, DK). Primary antibodies were rabbit monoclonal anti-cleaved Notch1 (Val1744) (Cell Signaling Technology, Leiden, NL) at 1:1000 and rat monoclonal anti-tubulin at 1:5000 (Abcam, Cambridge, UK).

### Immunostaining

All primary antibodies are listed with dilution in Supplementary Table 2. Dissected whole embryos (E10.5-E12.5) and foregut preparations (E13.5-E18.5) were fixed in 4% paraformaldehyde in PBS, embedded in Tissue-Tek O.C.T. (Sakura Finetek) and cryosectioned at 10 μm. For immunofluorescence analysis, antigen retrieval was conducted in pH6.0 citrate buffer, followed by permeabilization in 0.15% Triton X-100 in PBS. After blocking in 1% normal donkey serum in PBS with 0.1% Tween-20, sections were incubated overnight at 4°C with primary antibodies diluted in the same buffer. Primary antibodies were detected with anti-rabbit, guinea pig, mouse, rat, goat, sheep or chicken donkey-raised secondary antibodies conjugated to either Cy5 (1:500), Cy3 (1:1,000), Alexa Fluor 488 (1:1,000) or DyLight 405 (1:200) (all Jackson ImmunoResearch Europe, Ely, UK). Slides were mounted in Vectashield (Vector Laboratories, Burlingame, CA) with or without DAPI for counterstaining nuclei. Whole-mount IF of E10.5 whole embryos and E12.5, E15.5 and E18.5 foregut preparations was performed as previously described (Ahnfelt-Ronne et al., 2007). Specimens were cleared with BABB (benzyl alcohol:benzyl benzoate 1:2) then scanned confocally for *z*-stack image acquisition. Images were captured on a Leica SP8 or Zeiss LSM780 confocal microscope and figures prepared using Adobe Photoshop/Illustrator CS6 (Adobe Systems, San Jose, CA, USA). Cultured explants were fixed in 4% PFA for 40 min at room temperature followed by antigen retrieval for 1 hour at 37°C in citrate buffer (pH 6.0, 0.1 M 9.5% citric acid, 0.1 M 41.5% sodium citrate and 49% ddH_2_O). 3x 10 min wash in PBS at room temperature. Permeabilization in 1% Triton-X-100 diluted in PBS for 40 min. Block in 1% Normal Donkey Serum (#017-00-121) (Jackson ImmunoResearch Europe, Ely, UK) diluted in PBST. Explant were incubated in primary antibodies diluted in blocking buffer at 4°C overnight. The following day, explants were washed 3x 10 min in PSB followed by 2-hour incubation at room temperature with secondary antibodies diluted in blocking buffer.

### Antibody validation

Antisera against Dll1, Jag1, and Hes1 were validated by IF analysis of E10.5 neural tube from embryos that were either wildtype or null for the relevant gene (Figure S7). Two characteristic stripes of Jag1 in the neural tube (Johnston et al., 1997) and uniform but weak Jag1 in the E10.5 pancreas was detected by both anti-Jag1 antisera in wild type tissue, but not in equivalent *Jag1*-null tissue. The anti-Dll1 antibody detected Dll1 in the expected reciprocal pattern (compared to Jag1) in the neural tube and scattered cells in the pancreas of wild type embryos, but not *Dll1*^−/−^ embryos. The rabbit monoclonal anti-Hes1 antibody detected Hes1 in the expected patterns in both wild type tissues (i.e. prominently in floor plate and dorsal neural tube as well as pancreatic epithelium and weakly in the surrounding mesenchyme), but not in *Hes1*^−/−^ tissues.

### Bioluminescence imaging

Explants were imaged on a heated stage of an inverted microscope (Olympus IX83) and maintained at 37°C in 5% CO_2_. The bioluminescence signal was collected with an Olympus 40x UPLFN Universal Plan Fluorite oil immersion objective was transmitted directly to either a cooled Andor iXon Ultra 888 EMCCD camera or a cooled Andor iKon-M934 CCD camera (Oxford Instruments, Oxford, UK). The exposure time for Luc2-Hes1 was 5-10 min with no binning and for Dll1-Fluc it was 10 minutes with 4×4 binning.

### Bioluminescence quantification

Image sequences were analysed in Fiji (ImageJ version 2.0.0, NIH). Noise from cosmic rays was removed using the SpikeNoise Filter, while single-cell tracking was optimized using the Savitzky-Golay Temporal Filter in Fiji. To track single cells the path was defined with the ROI tool and supported by a Maximum Projection image. False colour (Fire) was used to illustrate the signal intensity in each cell. Z-axis Profiler Plus was used to extract the bioluminescence signal per cell over time for peak-to-peak quantification in Microsoft Excel. Mean amplitude for each oscillating cell was calculated as the difference in relative luciferase activity (a.u.) between individual peaks (P) and troughs (T) using the equation: Mean amplitude = (P_1_ – T_1_) + (P_2_ – T_2_) + … + (P_n_ – T_n_)/n. The local background arising from light scattering in the explants was measured in nine ROIs adjacent to oscillatory cells for each time-lapse movie and the average local background was subtracted from the raw bioluminescence signals before plotting tracks. Background from noise in the EMCCD camera was negligible.

### Cell quantification

~400 cells YFP^+^ lineage-traced cells were counted on 9-11 evenly spaced optical sections through the pancreas from each of three E12.5 N1IP::Cre^HI^; *Rosa26*^LSL-Ai3^ embryos and scored for co-expression of Sox9 and Ptf1a. YFP^+^ and GFP^+^ lineage-traced cells from *Hnf1b*-CreER^T2^; *R26*^YFP^ and *R26*^dnMaml1-GFP^ embryos, respectively, were counted on every fifth section throughout the pancreas, for a total of >200 cells/embryo, for each marker combination. Ptf1a^+^Nkx6.1^−^, Ptf1a^−^Nkx6.1^+^ and Ptf1a^+^Nkx6.1^+^ cells were quantified on every fifth section throughout dorsal pancreas from E12.5 controls (*R26*^Yfp/+^; *Sox17*^CreERT/+^), *Dll1*^ΔSox17Tam^, *Jag1*^ΔSox17Tam^, *Dll1*/*Jag1*^ΔSox17Tam^ using Imaris™ (Oxford Instruments, Oxford, UK). E12.5 *Jag1*^ΔFoxa2^ embryos were quantified the same way but with their own controls (see below). For quantification of E12.5 *Jag1*^ΔFoxa2^ Hes1 IF signal intensity in distal-most Ptf1a^+^ cells, corrected total cell fluorescence (CTCF) was determined using FIJI (Burgess et al., 2010; McCloy et al., 2014). Numbers of Ptf1a^+^Nkx6.1^−^, Ptf1a^−^Nkx6.1^+^ and Ptf1a^+^Nkx6.1^+^ cells as well as Sox9^+^ and insulin^+^ cells were manually scored from every tenth section of dorsal pancreata from E15.5 controls (*R26*^Yfp/+^; *Foxa2*^iCre/+^; *Dll1*^+/+^; *Jag1*^+/+^), *Jag1*^ΔFoxa2^, *Dll1*^ΔFoxa2^ and *Jag1/Dll1*^ΔFoxa2^ embryos and expressed relative to the area (in mm^2^) of the YFP^+^ dorsal pancreatic epithelium using Fiji.

### Statistical analyses

An F test was used to compare variances. Data sets with two groups having equal variances were analyzed by a two-tailed (unless otherwise indicated) Student’s t tests. For data with unequal variances we used two-tailed Welch’s t tests. Data sets with multiple groups were analyzed by one-way ANOVA, followed by Tukey’s post hoc test for multiple comparisons using the Prism statistical program (GraphPad, San Diego, CA). Results were expressed as mean ± S.D. and sample numbers are indicated in the figures. A minimum of three embryos per genotype were examined, except for *Jag1*; *Dll1*^ΔFoxa2^ for which three years of breeding have so far only yielded two embryos.

## Supplemental information

**Supplementary Table 1**

Genotyping primers.

D1VmC genotyping primers: Fwd: 5’-CTTCAAAGGACACCAAGTACCAGTCG-3’, WT-Rev: 5’-CTGTCCATAGTGCAATGGGAACAACC 3’, Venus-Rev: 5’ CTTGCTCACCATAAAGATGCGACCTCC 3’

J1VmC genotyping primers: Fwd: 5’ CAACACGGTCCCCATTAAGGATTACGAG 3’, WT-Rev: 5’ CTGTCCATAGTGCAATGGGAACAACC 3’, Venus-Rev: as above.

**Supplemental Table 2.** Antibodies employed in immunofluorescence analyses.

| Primary Antibodies |  |  |  |  |
| --- | --- | --- | --- | --- |
| Antigen | Species | Source | Catalogue # | Dilution |
| Pdx1 | Goat | NIH Beta Cell Biology Consortium (BCBC) | AB2027 | 1:10,000 |
| Sox9 | Guinea Pig | Gift from Ole Madsen/Novo Nordisk | N/A | 1:2,000 |
| Sox9 | Rabbit | EMD Millipore (Merck) | AB5535 | 1:1,000 |
| Ptf1a | Rabbit | NIH BCBC | AB2153 | 1:3,000 |
| Ptf1a | Guinea Pig | Gift from Jane E. Johnson, UT Southwestern Medical Center | N/A | 1:5,000 |
| CPA1 | Goat | R&D Systems | AF2765 | 1:200 |
| Amylase | Rabbit | Sigma-Aldrich (Merck) | A8273 | 1:500<sup>1</sup> |
| Mist1 | Rabbit mAb | Cell Signaling Technology | 14896 | 1:500<sup>1</sup> |
| Nkx6.1 | Mouse |  |  | 1:500<sup>2</sup> |
| Nkx6.1 | Rabbit | NIH BCBC | AB1069 | 1:2,000<sup>1</sup> |
| Ngn3 | Rabbit | NIH BCBC | AB2011 | 1:4,000 |
| Ngn3 | Chicken | NIH BCBC/Chris V. E. Wright | AB3854 | 1:1,000 |
| Chromogranin-A | Goat | Santa Cruz | sc-1488 | 1:200 |
| Insulin | Guinea Pig | Abcam | ab7842 | 1:200 |
| Insulin | Guinea Pig | Dako (Agilent) | A0564 | 1:800 |
| Glucagon | Mouse | Sigma-Aldrich (Merck) | G2654 | 1:800 |
| Glucagon | Guinea Pig | Millipore | 4031-01F | 1:4,000 |
| Somatostatin | Rabbit | Dako (Agilent) | A0566 | 1:2,000<sup>1</sup> |
| DBA | N/A | Vector Labs | B-1035 | 1:500<sup>3</sup> |
| Krt19 (CK19,<br>TROMA-III) | Rat | Developmental Studies Hybridoma Bank | N/A | 1:100<sup>4</sup> |
| Muc1 | Armenian<br>Hamster mAb | Thermo Fisher (Invitrogen) | MA5-11202 | 1:200 |
| ZO-1 (R26.4C) | Rat | Developmental Studies Hybridoma Bank | N/A | 1:1,000<sup>4</sup> |
| PKCζ | Rabbit | Santa Cruz | sc-216 | 1:800 |
| E-Cadherin | Mouse | BD Biosciences | 610181 | 1:2,000 |
| E-Cadherin | Rat | Novo Nordisk | N/A | 1:1,000 |
| Pecam1/CD31 | Rat | BD Biosciences | 550274 | 1:50 |
| Dll1 | Sheep | R&D Systems | AF3970 | 1:200 |
| Jag1 | Goat | Santa Cruz Biotechnology | sc-6011 | 1:200 |
| Jag1 | Rabbit mAb | Cell Signaling Technology | 2620 | 1:50 |
| Notch1 | Sheep | R&D Systems | AF5267 | 1:200 |
| Notch2 | Goat | R&D Systems | AF1190 | 1:200 |
| Hes1 | Rabbit mAb | Cell Signaling Technology | 11988 | 1:200 |
| GFP | Chicken | Abcam | ab13970 | 1:1,000 |
| GFP | Rabbit | Clontech | 632460 | 1:1,000 |
| RFP | Rabbit | Rockland | 600-401-379 | 1:2,000 |

<sup>1</sup> Detected with (1:500) Cy3-conjugated F(ab')<sub>2</sub> fragment donkey anti-rabbit IgG (Jackson
ImmunoResearch Europe; 711-166-152)
<sup>2</sup> Blocking and antibody incubations performed using M.O.M. Basic Kit (Vector Labs; BMK-
<sup>3</sup> Biotinylated DBA detected with (30 µg/ml) AMCA Avidin D (Vector Labs; A-2008)
<sup>4</sup> Concentrate

**Movie S1.** Time-lapse movie of explanted E10.5 Luc2-Hes1 dorsal pancreatic bud cultured for 24 hours before imaging commenced. Individual Luc2-Hes1 expressing nuclei are displaying ultradian oscillations. The movie shows 20 hours of development and each frame is 10 minutes.

**Movie S2.** Time-lapse movie of explanted E10.5 Luc2-Hes1 dorsal pancreatic bud cultured for 96 hours before imaging commenced. Individual Luc2-Hes1 expressing nuclei are displaying ultradian oscillations. The movie shows 20 hours of development and each frame is 10 minutes.

**Movie S3.** Time-lapse movie of explanted E10.5 Luc2-Hes1 dorsal pancreatic bud cultured for 144 hours before imaging commenced. Individual Luc2-Hes1 expressing nuclei are displaying ultradian oscillations. The movie shows 20 hours of development and each frame is 10 minutes.

**Movie S4.** Time-lapse movie of explanted E12.5 Dll1-Fluc dorsal pancreatic bud cultured for 4 hours before imaging commenced. Individual Luc2-Hes1 expressing nuclei are displaying ultradian oscillations. The movie shows 20 hours of development and each frame is 10 minutes.

