## Supplemental Figures for "Jag1 modulates an oscillatory Dll1-Notch-Hes1 signaling module to coordinate growth and fate of pancreatic progenitors"

**Supplemental Figure legends**

**Figure S1.** Expression of Jag1 and Dll1 in the developing pancreas.

(A) Sections of E9.5, E10.5, E11.5, E12.5 and E15.5 *Jag1*$^{J1VmC}$ and *Dll1*$^{D1VmC}$ embryos stained for mCherry, Pdx1 and Sox9 as indicated. Arrows: Proximal mCherry$^{-}$ cells. Arrowheads: Peripheral mCherry$^{+}$ cells Higher magnifications of boxed areas are shown in insets. dp: dorsal pancreas; vp: ventral pancreas; bp: bi-potent progenitor domain; ac: emerging acini; v: vessel. Scale bar is 20 µm.

(B) Sections of E10.5, E12.5, and E15.5 dorsal pancreata stained for Jag1 in combination with either Pecam and Cdh1, Ngn3 and Gcg, or Ptf1a and Sox9 as indicated. Arrowheads: Expression of Jag1 in Ptf1a$^{+}$ PACs in the distal E12.5 epithelium. bp: bi-potent progenitor domain; ec: endocrine cells; ac: emerging acini; v: vessel. Scale bars are 25 µm.

(C) Sections of E10.5, E12.5, and E15.5 dorsal pancreata stained for Dll1 in combination with either Pecam and Cdh1, Ngn3 and Gcg, or Ptf1a and Sox9 as indicated. Arrows: Expression ofDll1 in Ptf1a$^{+}$ MPCs at E10.5, PACs in the distal E12.5 epithelium, and emerging acini at E15.5. Arrowheads: Dll1 expression in Ngn3$^{+}$ cells at all stages and in Sox9$^{+}$ BPs at E12.5 and E15.5.

(D) Sections of E9.5, E10.5, E11.5, E12.5 and E15.5 *Jag1*$^{J1VmC}$; *Hes1*-EGFP and *Dll1*$^{D1VmC}$; *Hes1*-EGFP embryos stained for mCherry, EGFP, and Sox9 as indicated. Arrowheads in *Jag1*$^{J1VmC}$ panels: mCherry co-expression with EGFP in most E10.5 MPCs (Sox9$^{+}$) but mCherry segregated to distal EGFP$^{-}$ cells through E11.5 and E12.5. Arrows in *Jag1*$^{J1VmC}$ panels: EGFP$^{+}$ proximal cells from E11.5-E15.5. Arrowheads in *Dll1*$^{D1VmC}$ panels: Heterogeneous mCherry frequently co-expressed with EGFP in E9.5-E10.5 MPCs and in proximal BPs from E11.5-E15.5. Arrows in *Dll1*$^{D1VmC}$ panels: mCherry single positive cells in the distal region throughout development. dp: dorsal pancreas; vp: ventral pancreas; bp: bi-potent progenitor domain; ac: emerging acini.

(E) Schematic representation of the expression patterns of Notch ligands and receptors at E10.5 and E12.5. Yellow indicates Jag1-Dll1 co-expression. Lighter color shades indicate lower expression levels. ec: endocrine clusters.

**Figure S2.** Stage-dependent allocation of progenitor fate by Notch suppression.

(A) Overview of strategy applied to identify fates of progeny from pancreatic progenitors. Approximate temporal windows of Tamoxifen (Tam) activity resulting from single intraperitoneal injections at E11.5, E12.5 or E13.5 and cell fate-specific markers are indicated. PAC: pre-acinar cell; BP: bipotent progenitor; EP: endocrine precursor; EC: endocrine cell.

(B-M) Sections of E15.5 *Sox9*-CreER$^{T2}$; *R26*$^{YFP}$ control (B-D, H-J) or *Sox9*-CreER$^{T2}$; *R26*$^{dnMaml1}$ Notch-blocked pancreata (E-G, K-M) stained for EYFP/EGFP and Sox9 combined with either Ptf1a (B-G) or Ngn3 and Chga (H-M) as indicated. bp: bi-potent progenitor domain; ac: emerging acini; ec: endocrine cells. Scale bar 50 μm. Arrows in (B-D): Lineage-labeled Sox9$^{+}$ BPs. Arrowheads in (B-G): Lineage-labeled Ptf1a$^{+}$ PACs. Arrows in (F-G): Lineage-labeled Ptf1a/Sox9 double negative cells. Arrows in (H-J): Lineage-labelled Sox9$^{+}$ BPs. Arrowheads in (I-J): Lineage-labelled Ngn3$^{+}$ endocrine precursors**.** Arrowheads in (K-M): Lineage-labeled Chga$^{+}$ endocrine cells. Arrows in (K): Lineage-labeled Ngn3/Chga/Sox9 triple negative cells.

**Figure S3.** Delayed resolution of PD patterning in the *Jag1*$^{\Delta Foxa2}$ pancreas.

(A-C) Sections of E13.5 and E14.5 pancreata from *Jag1*$^{\Delta Foxa2/+}$ and *Jag1*$^{\Delta Foxa2/-}$ embryos stained for Ptf1a and Nkx6-1 (A), Ptf1a and Sox9 (B), or insulin (Ins) and Ngn3 (C) as indicated. Insets show higher magnifications of individual channels. Scale bars in main panels are 50 μm and 25 μm in insets.

**Figure S4.** Heterozygote phenotype in E15.5 *Jag1*$^{\Delta Foxa2}$ pancreas.

(A) IF for Ptf1a and Nkx6-1 on E15.5 *Jag1*$^{\Delta Foxa2/+}$ and *Jag1*$^{\Delta Foxa2/-}$ pancreata. Insets show higher magnifications of distal (Di) and proximal (Pr) regions.

(B) Quantitatively, Nkx6-1$^{+}$ cells are progressively lost while Ptf1a$^{+}$ are proportionally increased in E15.5 *Jag1*$^{\Delta Foxa2/+}$ and *Jag1*$^{\Delta Foxa2/-}$ pancreata compared to wildtype controls (mean ± S.D., N = 3).

(C) IF for Sox9, Ngn3 and insulin on E15.5 *Jag1*$^{\Delta Foxa2/+}$ and *Jag1*$^{\Delta Foxa2/-}$ pancreata. Insets show higher magnifications of distal (Di) and proximal (Pr) regions. The pancreatic epithelium is demarcated by dashed lines.

(D) Quantitatively, BPs and β-cells are progressively reduced in E15.5 *Jag1*$^{\Delta Foxa2/+}$ and *Jag1*$^{\Delta Foxa2/-}$ pancreata compared to wildtype controls (mean ± S.D., N = 3). Scale bars,

100 μm (main panels) and 25 μm (insets). Note that wildtype and homozygous mutant data in A-D are the same as shown in Figure 5.

(E) Sections of E15.5 *Jag1*$^{\Delta Foxa2/+}$ and *Jag1*$^{\Delta Foxa2/-}$ pancreata stained for Ptf1a, Sox9 and Muc1. Note that Sox9$^{+}$ terminal duct cells connecting to acini in controls are replaced by Ptf1a$^{+}$ elongated "duct"-like structures in *Jag1*$^{\Delta Foxa2/-}$ mutants in both proximal (Pr) and distal (Di) regions. Boxed areas are shown at higher magnification in insets.

(F) Sections of E15.5 *Jag1*$^{\Delta Foxa2/+}$ and *Jag1*$^{\Delta Foxa2/-}$ pancreata stained for Ptf1a, Mist1 and Pdx1. Scale bars are 100 μm (main panels) and 25 μm (insets).

**Figure S5.** E15.5 *Mib1*$^{\Delta Ptf1a}$ pancreas lacks CAC precursors.

(A-B) Sections of E15.5 *Mib1*$^{\Delta Ptf1a/+}$; *R26*$^{YFP/+}$ (A) and *Mib1*$^{\Delta Ptf1a/\Delta Ptf1a}$; *R26*$^{YFP/+}$ (B) pancreata stained for Ptf1a, Sox9 and YFP as indicated. Insets show higher magnification views of boxed areas in Pr (proximal) and Di (distal) regions. Arrows in Pr insets: recombination is mosaic in the Sox9$^{+}$ domain but uniformly high in the Ptf1a$^{+}$ domain. Arrows in Di inset: The terminal most Sox9$^{+}$ cells located within forming acini of the *Mib1*$^{\Delta Ptf1a/+}$ pancreas are missing from the *Mib1*$^{\Delta Ptf1a/\Delta Ptf1a}$ pancreas (asterisk). Scale bars are 100 μm (main panels) and 20 μm (insets).

**Figure S6.** Model for Notch pathway function in MPC maintenance and fate choice.

(A) MPC maintenance and growth: Initially, MPCs proliferate at a given rate, in part determined by input from Notch receptors. High levels of the ligand (L) Dll1 in one cell mediate receptor (R) activation in neighbors resulting in activation of Hes1. A low level of Jag1 attenuates receptor activation by sequestering a fraction of the receptors (R *cis*-inhibition). Ultradian oscillations in Hes1 and Dll1 enable a temporal symmetry where MPCs alternate between sending and receiving mitogenic input via Notch. Deletion of Dll1 lowers peak ligand levels drastically, leading to a state where surplus receptors *cis*-inhibit the remaining Jag1 ligands. As a result, NICD-mediated Hes1 activation (amplitude) is suppressed and mitogenic input is reduced. In contrast, deletion of Jag1 removes its dampening effect on Notch activity resulting in increased Dll1-mediated signaling and MPC proliferation but does not affect oscillations. Stable intermediate levels of Dll1 in *Dll1*$^{T1/T1}$ and *Dll1*$^{T2/T2}$ mutants causes stable receptor *cis*-

inhibition and prevent cells from entering a Dll1$^{Lo}$ state required for signal reception. The outcome is decreased proliferation, similar to loss of Dll1 altogether.

(B) Increased Jag1 expression breaks the symmetry by sequestering all receptors in emerging Dll1$^{Hi}$Jag1$^{Hi}$ pro-acinar cells (PACs) and bolstering free Jag1 and Dll1 levels. As a result, *trans*-activation is augmented in nascent bi-potent progenitors (BPs), which is further reinforced by onset of Notch2 expression and loss of Jag1-mediated attenuation. Deletion of Dll1 causes a slight bias towards PAC fate but high levels of Jag1 allow MPCs to exit the multipotent state normally. In contrast, loss of Jag1 prevents symmetry breaking and exit from the multipotent state, except in the central-most progenitors, which receive high Dll1 input from Ngn3$^{+}$ endocrine precursors.

(C) The genetic circuitry (arrows) and protein-protein interactions (double line arrows) underlying the regulatory logic determining cell fate choices. Black indicates active, and grey inactive, components and connections. The Ptf1a/Rbpj complex (Ptf1) activates *Dll1* expression while Hes1 represses *Ptf1a*, *Dll1* and itself, with the latter likely causing the observed ultradian oscillations of both Hes1 and Dll1. The points of action for DAPT and MLN4924 are indicated.

**Figure S7.** Validation of anti-Dll1, -Jag1 and -Hes1 antisera.

(A-D) Rabbit monoclonal anti-Jag1 antibody (Cell Signaling Technology, #2620) gives a membranous signal on cells in the neural tube (A-B) and a weaker but specific membranous staining throughout the dorsal (dp) and ventral (vp) pancreatic buds (C-D) in E10.5 control mice (C) but not *Jag1*$^{-/-}$ littermates (D).

(E-H) Similarly, goat anti-Jag1 antibody (Santa Cruz Biotechnology, #sc-6011) gives membranous signal on cells in the neural tube (E-F) and weaker but specific membranous staining throughout both pancreatic buds (G-H) in E10.5 control embryos (E, G) but not *Jag1*$^{-/-}$ littermates (F, H).

(I-L) Sheep anti-Dll1 antibody (R&D Systems, #AF3970) yields heterogeneous membranous signal on cells in the neural tube (I) and weaker but specific signal in both pancreatic buds (K) in E10.5 control embryos but not nullizygous *Dll1*$^{lacZ/lacZ}$ littermates (J, L).

(M-P) Rabbit monoclonal anti-Hes1 antibody (Cell Signaling Technology, #11988) yields strong nuclear Hes1 signal in the dorsal neural tube and floorplate (fp) (M-N) and throughout both pancreatic buds (O-P) in E10.25 control embryos (M, O) but not *Hes1*$^{-/-}$ littermates (N, P). Scale bars are 50 μm. Sox9 signal is shown for reference. Weak signals in pancreatic buds compared to neural tube are visualized by enhancing the brightness equally for control and nullizygous images in Adobe Photoshop. The pancreatic epithelium is demarcated by dashed lines where necessary.

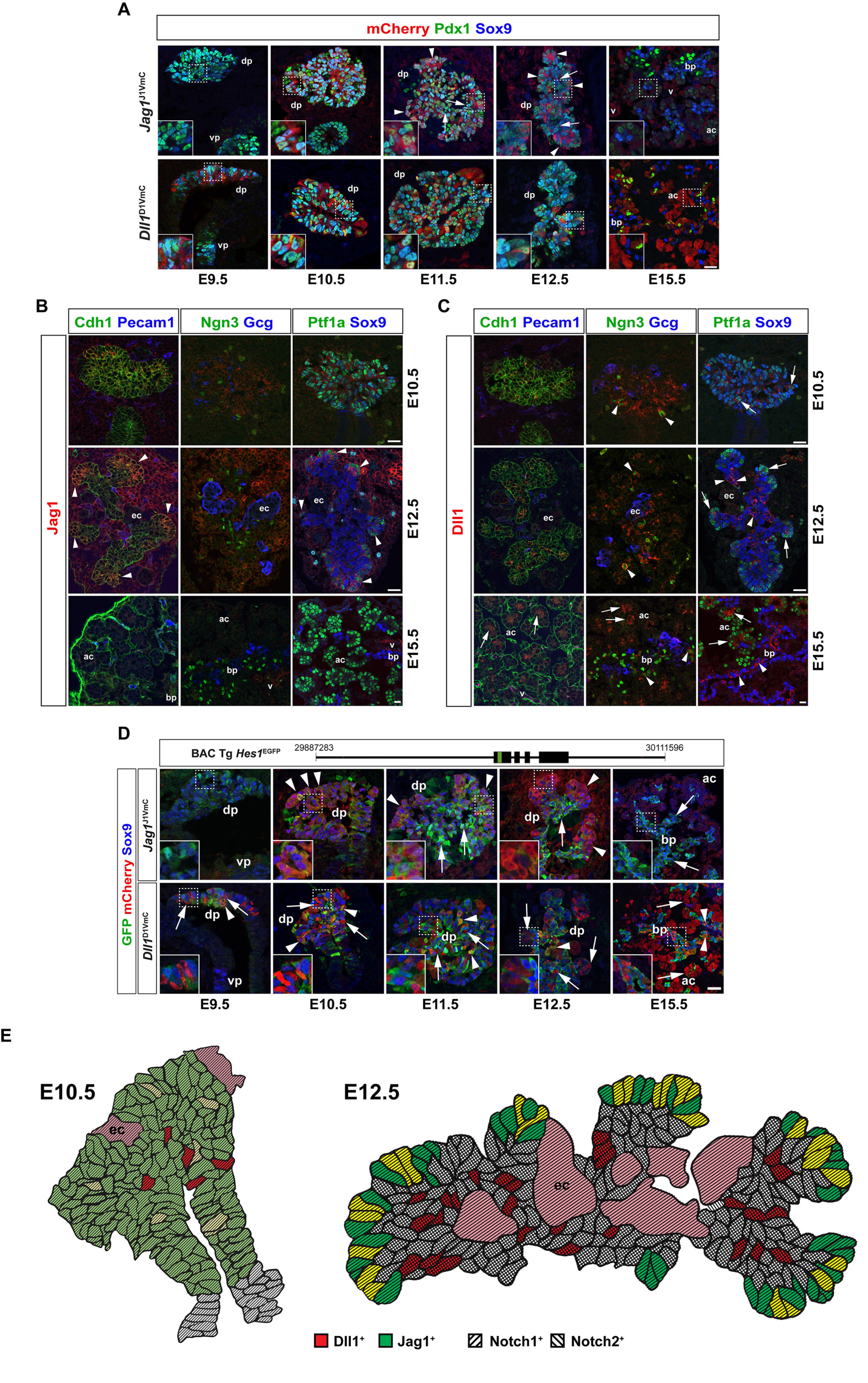

A
mCherry Pdx1 Sox9
Jag1J1VmC
Dll1D1VmC
dp
vp
bp
v
ac
E9.5
E10.5
E11.5
E12.5
E15.5
B
Cdh1 Pecam1
Ngn3 Gcg
Ptf1a Sox9
Jag1
ec
C
Dll1
D
BAC Tg Hes1EGFP
29887283
30111596
GFP mCherry Sox9
E
E10.5
E12.5
Dll1+
Jag1+
Notch1+
Notch2+


**Figure S1**

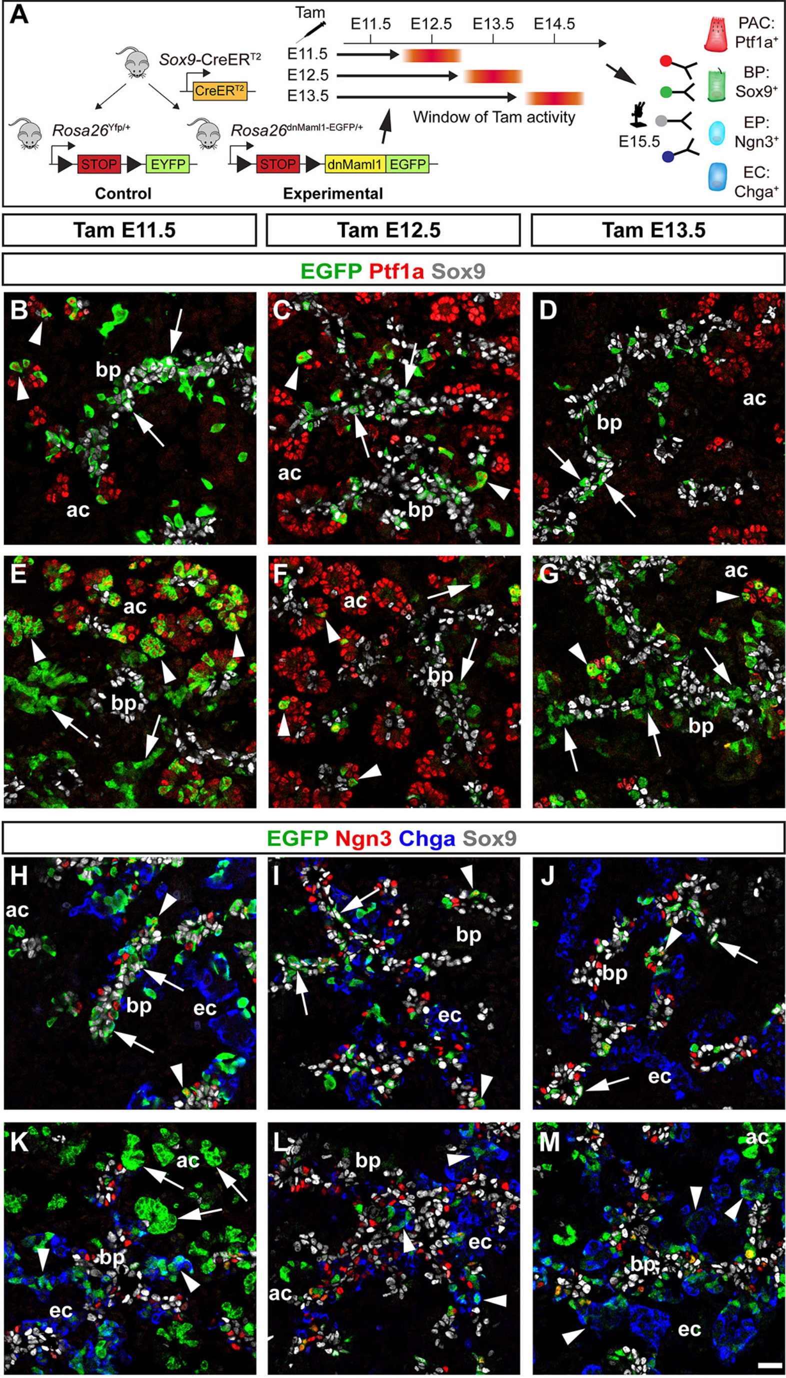
A
Tam
E11.5
E12.5
E13.5
E14.5
Sox9-CreERT2
CreERT2
Rosa26Yfp/+
Rosa26dnMaml1-EGFP/+
STOP
EYFP
dnMaml1
EGFP
Control
Experimental
Window of Tam activity
E15.5
PAC: Ptf1a+
BP: Sox9+
EP: Ngn3+
EC: Chga+
Tam E11.5
Tam E12.5
Tam E13.5
EGFP Ptf1a Sox9
R26YFP
R26dnMaml1
EGFP Ngn3 Chga Sox9
B
C
D
E
F
G
H
I
J
K
L
M
bp
ac
ec

Figure S2

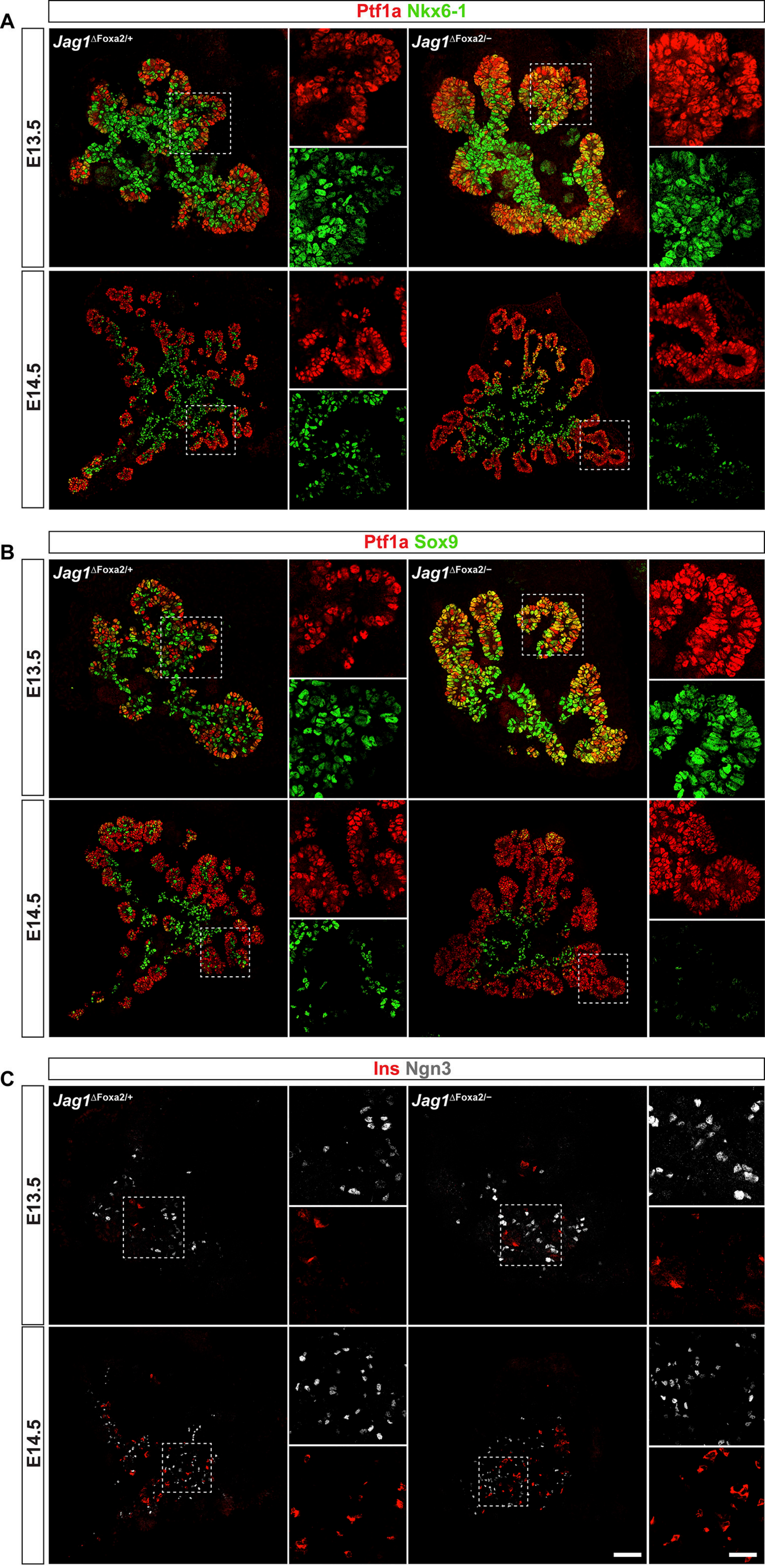
A
Ptf1a Nkx6-1
Jag1ΔFoxa2/+
Jag1ΔFoxa2/−
E13.5
E14.5
B
Ptf1a Sox9
Jag1ΔFoxa2/+
Jag1ΔFoxa2/−
E13.5
E14.5
C
Ins Ngn3
Jag1ΔFoxa2/+
Jag1ΔFoxa2/−
E13.5
E14.5

**Figure S3**

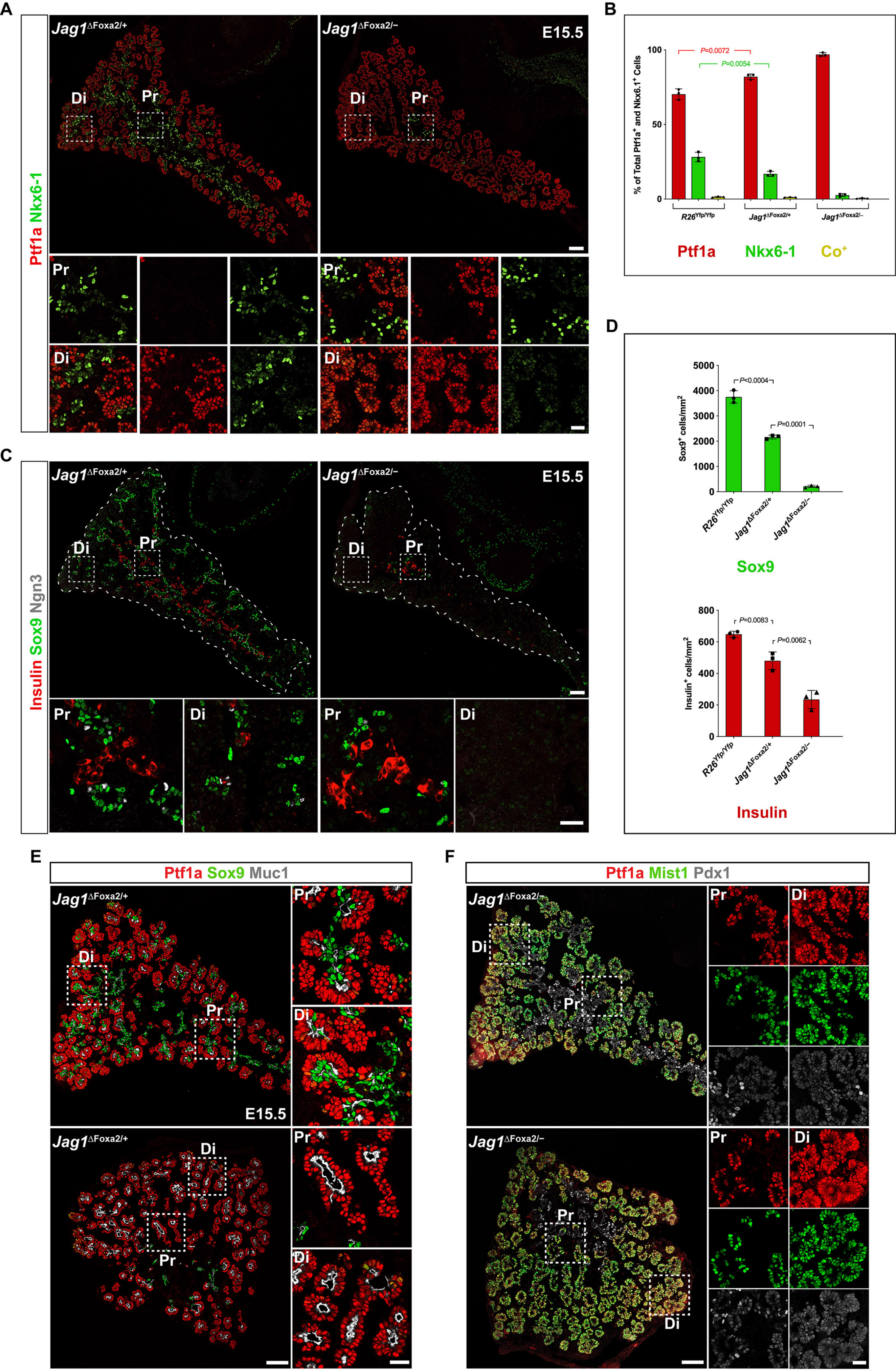
A
B
C
D
E
F
Ptf1a Nkx6-1
Jag1ΔFoxa2/+
Jag1ΔFoxa2/–
E15.5
Di
Pr
% of Total Ptf1a+ and Nkx6.1+ Cells
P=0.0072
P=0.0054
R26Yfp/Yfp
Ptf1a
Nkx6-1
Co+
Insulin Sox9 Ngn3
Sox9+ cells/mm2
P<0.0004
P=0.0001
Sox9
Insulin+ cells/mm2
P=0.0083
P=0.0062
Insulin
Ptf1a Sox9 Muc1
Ptf1a Mist1 Pdx1

Figure S4

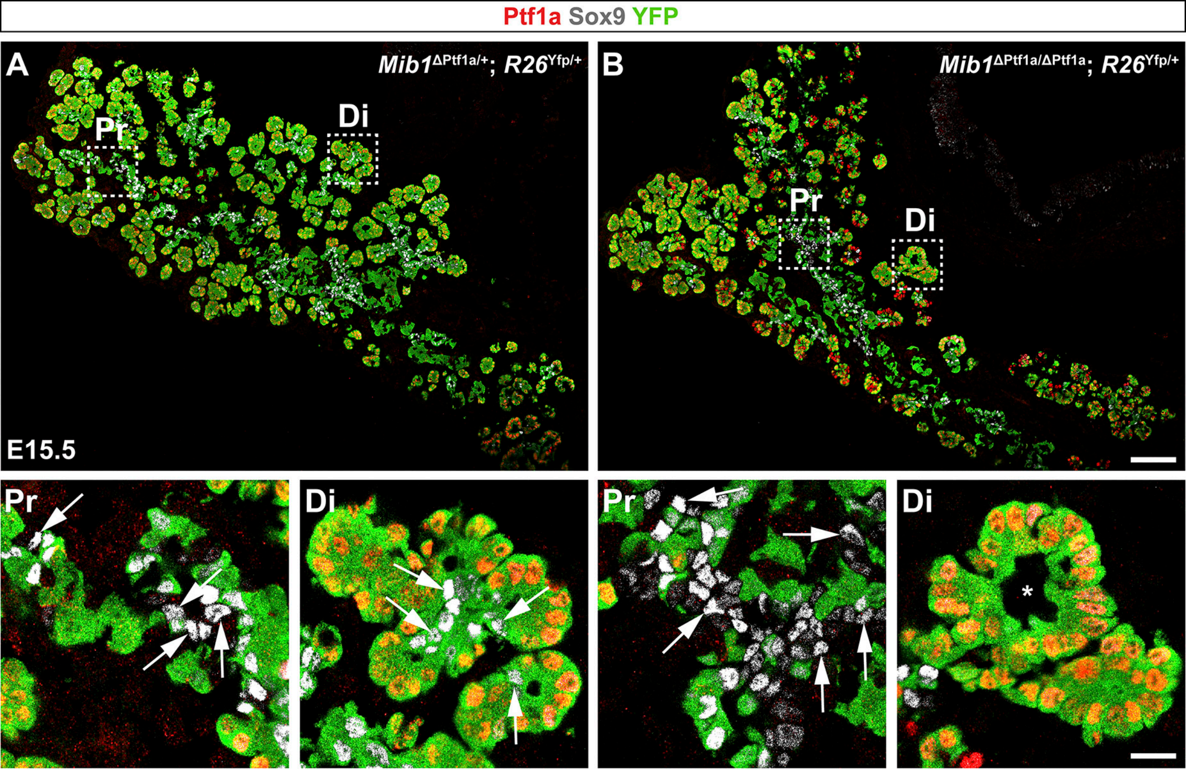
Ptf1a Sox9 YFP
A
Mib1ΔPtf1a/+; R26Yfp/+
Pr
Di
E15.5
B
Mib1ΔPtf1a/ΔPtf1a; R26Yfp/+
Pr
Di
Pr
Di
Pr
Di

**Figure S5**

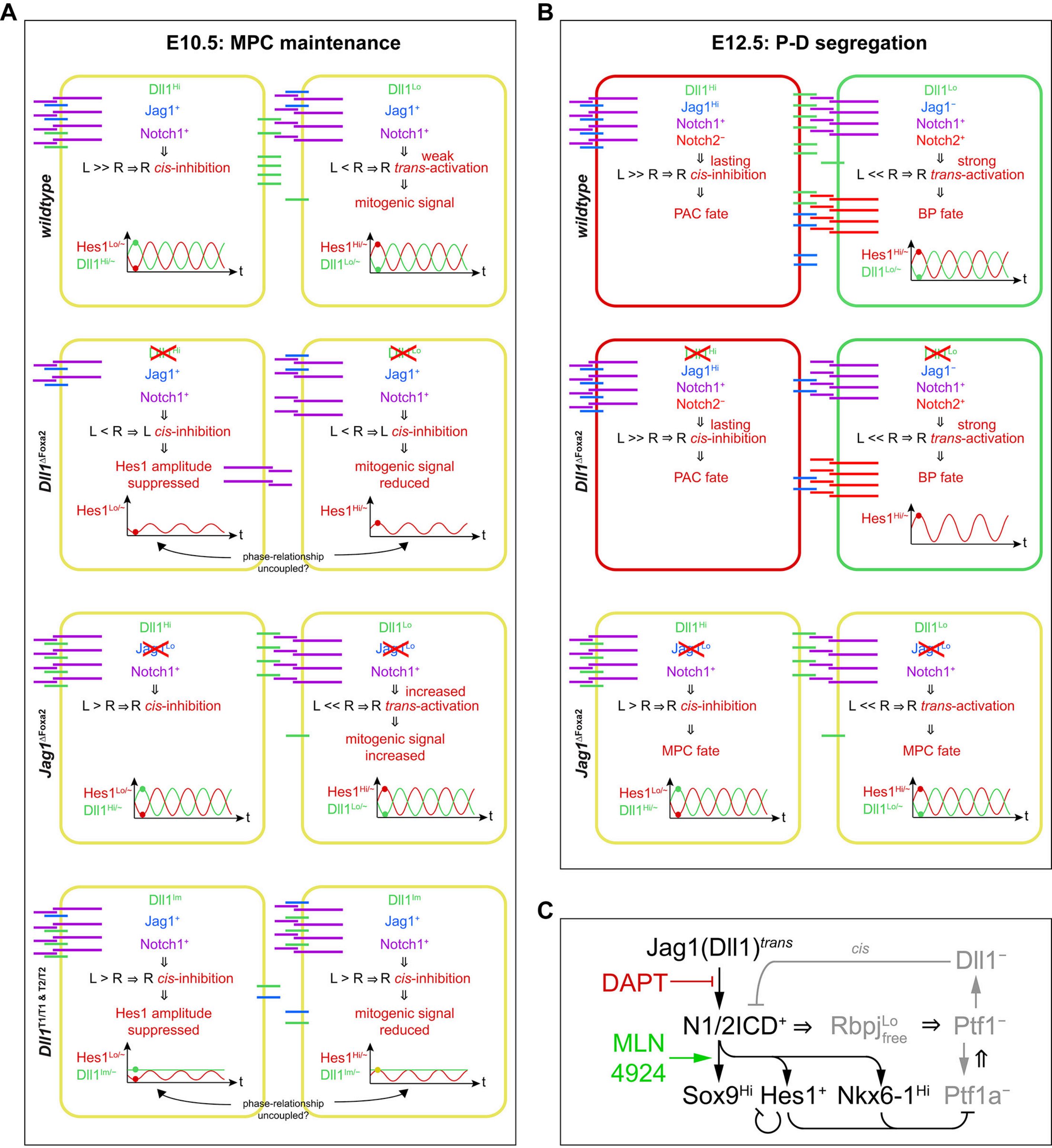
A
E10.5: MPC maintenance
wildtype
Dll1$^{Hi}$
Jag1$^{+}$
Notch1$^{+}$
L >> R ⇒ R cis-inhibition
Hes1$^{Lo/\sim}$
Dll1$^{Hi/\sim}$
t
Dll1$^{Lo}$
Jag1$^{+}$
Notch1$^{+}$
weak
L < R ⇒ R trans-activation
mitogenic signal
Hes1$^{Hi/\sim}$
Dll1$^{Lo/\sim}$
Dll1$^{\Delta Foxa2}$
L < R ⇒ L cis-inhibition
Hes1 amplitude suppressed
mitogenic signal reduced
phase-relationship uncoupled?
Jag1$^{\Delta Foxa2}$
Jag1$^{Lo}$
L > R ⇒ R cis-inhibition
increased
L << R ⇒ R trans-activation
mitogenic signal increased
Dll1$^{T1/T1 \& T2/T2}$
Dll1$^{Im}$
L > R ⇒ R cis-inhibition
Hes1 amplitude suppressed
mitogenic signal reduced
Dll1$^{Im/\sim}$
B
E12.5: P-D segregation
Jag1$^{Hi}$
Notch2$^{-}$
lasting
L >> R ⇒ R cis-inhibition
PAC fate
Jag1$^{-}$
Notch2$^{+}$
strong
L << R ⇒ R trans-activation
BP fate
MPC fate
C
Jag1(Dll1)$^{trans}$
cis
Dll1$^{-}$
DAPT
N1/2ICD$^{+}$ ⇒ Rbpj$^{Lo}_{free}$ ⇒ Ptf1$^{-}$
MLN 4924
Sox9$^{Hi}$ Hes1$^{+}$ Nkx6-1$^{Hi}$ Ptf1a$^{-}$

**Figure S6**

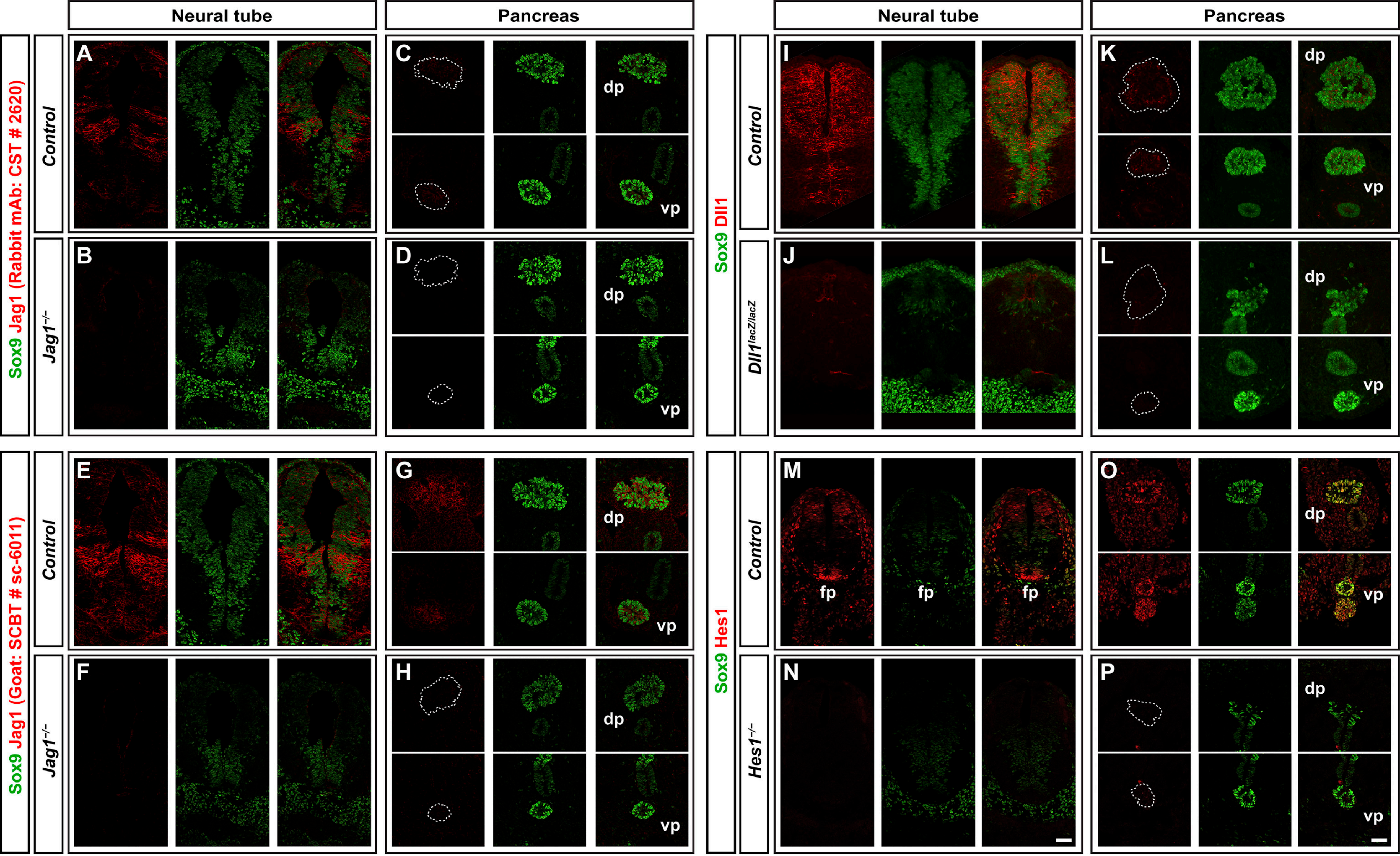

Neural tube
Pancreas
Neural tube
Pancreas
Sox9 Jag1 (Rabbit mAb: CST # 2620)
Control
Jag1−/−
A
B
C
D
dp
vp
Sox9 Dll1
Control
Dll1lacZ/lacZ
I
J
K
L
Sox9 Jag1 (Goat: SCBT # sc-6011)
E
F
G
H
Sox9 Hes1
Hes1−/−
M
N
O
P
fp


**Figure S7**
